# Targeting the PI5P4K lipid kinase family in cancer using novel covalent inhibitors

**DOI:** 10.1101/819961

**Authors:** Sindhu Carmen Sivakumaren, Hyeseok Shim, Tinghu Zhang, Fleur M. Ferguson, Mark R. Lundquist, Christopher M. Browne, Hyuk-Soo Seo, Marcia N. Paddock, Theresa D. Manz, Baishan Jiang, Ming-Feng Hao, Pranav Krishnan, Diana G. Wang, T. Jonathan Yang, Nicholas P. Kwiatkowski, Scott B. Ficarro, James M. Cunningham, Jarrod A. Marto, Sirano Dhe-Paganon, Lewis C. Cantley, Nathanael S. Gray

## Abstract

The PI5P4Ks have been demonstrated to be important for cancer cell proliferation and other diseases. However, the therapeutic potential of targeting these kinases is understudied due to a lack of potent, specific small molecules available. Here we present the discovery and characterization of a novel pan-PI5P4K inhibitor, THZ-P1-2, that covalently targets cysteines on a disordered loop in PI5P4Kα/β/γ. THZ-P1-2 demonstrates cellular on-target engagement with limited off-targets across the kinome. AML/ALL cell lines were sensitive to THZ-P1-2, consistent with PI5P4K’s reported role in leukemogenesis. THZ-P1-2 causes autophagosome clearance defects and upregulation in TFEB nuclear localization and target genes, disrupting autophagy in a covalent-dependent manner and phenocopying the effects of PI5P4K genetic deletion. Our studies demonstrate that PI5P4Ks are tractable targets, with THZ-P1-2 as a useful tool to further interrogate the therapeutic potential of PI5P4K inhibition and inform drug discovery campaigns for these lipid kinases in cancer metabolism and other autophagy-dependent disorders.

## Introduction

Phosphatidylinositol 5-phosphate (PI-5-P) is one of the seven phosphosinositides that regulate a wide range of cellular functions (Balla, Szentpetery, & Kim, 2009; Bulley, Clarke, Droubi, Giudici, & Irvine, 2015). Known to localize in the nucleus, Golgi, endoplasmic reticulum and at the plasma membrane, PI-5-P is an oxidative stress-induced regulator of AKT activation and is also regulated by proline isomerase Pin1 through Pin1’s interactions with the phosphatidylinositol 5-phosphate 4-kinases (PI5P4K) in times of cellular stress (Keune et al., 2012; Keune et al., 2013). The phosphorylation of the low abundance phosphoinositide PI-5-P at the 4-position, yielding the product phosphatidylinositol-4,5-bisphosphate (PI-4,5-P_2_), is catalyzed by PI5P4Kα, β and γ (encoded by genes *PIP4K2A, PIP4K2B* and *PIP4K2C*) (Rameh et al., 1997; Rameh & Cantley, 1999). PI-4,5-P_2_ is an important precursor for second messengers inositol-1,4,5-triphosphate (IP3), diacylgycerol (DAG) and phosphatidylinositol-3,4,5-trisphosphate (PI-3,4,5-P_3_) (Martelli et al., 1992; Divecha et al., 1993; Fiume et al., 2012; Fiume et al., 2015).

While the majority of PI-4,5-P_2_ is generated by phosphorylation of phosphatidylinositol 4-phosphate (PI-4-P) on the 5-position by the Type I PI4P5K kinases, a PI5P4K-driven alternate route was discovered in 1997, hence the designation Type II (Rameh et al., 1997). The PI5P4Ks were traditionally thought to mainly be crucial direct regulators of PI-5-P levels (Bulley et al., 2015; Stijf-Bultsma & Sommer et al., 2015; Hasegawa, Strunk, & Weisman, 2017) however, PI5P4Kα was found to synthesize a pool of PI-4,5-P_2_ that is specifically important in mTORC2 regulation (Bulley et al., 2016) and to play a critical role in intracellular cholesterol transport by modulating PI-4,5-P_2_ homeostasis on peroxisome membranes (Hu et al., 2018). The low-activity isoform PI5P4Kγ was demonstrated to positively regulate Notch1 signaling by facilitating receptor recycling, suggesting that endosome-localized production of PI(4,5)P_2_ is involved Notch transport (Zheng & Conner, 2018). PI5P4Kα/β were also shown to be required for autophagosome-lysosome fusion during times of metabolic stress, suggesting that they were evolved by multicellular organisms to produce sufficient PI-4,5-P_2_ in nutrient-deficient conditions (Lundquist et al., 2018). These findings have dispelled the notion of PI5P4K as simply being functionally redundant in PI-4,5-P_2_ production.

PI5P4K has been suggested to be important in several diseases. *PIP4K2B*-null mice were found to have increased insulin sensitivity and reduced adiposity (Lamia et al., 2004). *PIP4K2A* was found to be a dependency in AML and ALL (Jude et al., 2015; Rosales-Rodríguez, et al., 2016; Urayama et al., 2018) and *PIP4K2A*^*-/-*^, *PIP4K2B*^+/-^, *TP53*^*-/-*^ mice had a dramatic tumor-free life extension compared to *TP53*^*-/-*^ mice, uncovering a potential synthetic lethality of PI5P4K with p53 (Emerling et al., 2013). Knockdown of *PIP4K2A/B* in human retinal pigment epithelial cells and rabbit models abrogated the pathogenesis of proliferative vitreoretinopathy (Ma et al., 2016). Deletion of *PIP4K2C* in mice resulted in an increase of proinflammatory cytokines and T-helper-cells, as well as a decrease in regulatory T-cells via hyperactivation of mTORC1 signaling (Shim et al., 2016). Pharmacological inhibition or knockdown of PI5P4Kγ reduced mutant huntingtin protein in human patient fibroblasts and aggregates in neurons, and relieved neuronal degeneration in *Drosophila* models of Huntington’s disease (Al-Ramahi et al., 2017). The critical role of the PI5P4Ks in mediating autophagy may explain their induced essentiality in various disease pathologies (Emerling et al., 2013; Vicinanza et al., 2015; Al-Ramahi et al., 2017; Lundquist et al., 2018). Collectively, these studies suggest that the PI5P4Ks represent a novel lipid kinase family whose underlying biology is important to numerous cellular processes and warrants further investigation of their therapeutic potential across a range of disease states.

The relevance of PI5P4K in a wide range of diseases has motivated efforts to develop PI5P4K inhibitors. Reported pan-PI5P4K inhibitors (Kitagawa et al., 2017) and isoform-specific inhibitors of PI5P4Kα (Davis et al., 2013), PI5P4Kβ (Voss et al., 2014) and PI5P4Kγ (Clarke et al., 2015; Al-Ramahi et al., 2017) have laid the foundation for evidence of PI5P4K druggability and motivated a need for inhibitors with further improved pharmacological properties. Here we present the identification of a novel potent, covalent PI5P4K inhibitor, THZ-P1-2, and characterize its cellular pharmacology in the contexts of autophagy and cancer. Using a multipronged approach combining biochemical and cellular assays, mass spectrometry, and crystallography, we discovered that THZ-P1-2 inhibits the PI5P4K family at sub-micromolar concentrations by reacting covalently with cysteine residues in a flexible loop outside the kinase domain of all three kinase isoforms. We show that THZ-P1-2 exhibits a reasonable selectivity profile across the kinome, with an S-score S_(10)_ of 0.02 (Karaman et al., 2008, Davis et al., 2011) and inhibits cell proliferation at micromolar concentrations in a panel of leukemia cell lines in a manner dependent on covalent targeting. Finally, further investigation of the mechanism of action of THZ-P1-2 led us to the observation that THZ-P1-2 causes lysosomal disruption and defects in the clearance of autophagosomes, halting autophagy and phenocopying the effects of genetic deletion of PI5P4Kα/β/γ (Al-Ramahi et al., 2017; Lundquist et al., 2018). These studies provide evidence that irreversible inhibition of PI5P4K by THZ-P1-2 compromises autophagy, an essential alternative energy source during periods of metabolic stress which cancer cells depend on to maintain cellular homeostasis and prolonged cell viability, further suggesting PI5P4K as a potentially important target in cancer metabolism and other autophagy-dependent diseases.

## Results

### Chemoproteomic profiling and synthetic chemistry approaches reveal novel lead PI5P4K inhibitor THZ-P1-2

A previously developed acrylamide-based covalent JNK inhibitor, JNK-IN-7, was serendipitously discovered to also have activity on PI5P4Kγ through KiNativ cellular selectivity profiling (Fig. 1A, Fig. S1A) (Zhang et al., 2012). JNK-IN-7 was found to inhibit the kinase activity of PI5P4Kα/β/γ in a radiometric thin-layer chromatography (TLC) assay (Fig. S1B-C) and mass spectrometry revealed labeling of cysteine residues located on a disordered loop that is not observed in the available PI5P4K crystal structures (Fig. S1D-E). We pursued a focused medicinal chemistry campaign guided by biochemical kinase assays and cellular pulldowns to optimize the potency and selectivity of phenylamino-pyrimidine which resulted in the development of THZ-P1-2 (Fig. 1A) (full structure-activity relationships to be described elsewhere). THZ-P1-2 demonstrated inhibition of PI5P4Kα kinase activity (Fig. 1B), with an IC_50_ of 190 nM using an ADP-Glo (Promega) bioluminescence assay. THZ-P1-2 exhibited approximately 75% inhibition of PI-4,5-P_2_ formation by PI5P4Kα and PI5P4K γ and 50% inhibition by PI5P4Kβ at a concentration of 0.7 μM monitored using a thin-layer chromatography (TLC) assay (Fig. 1C-D).

**Figure 1.**
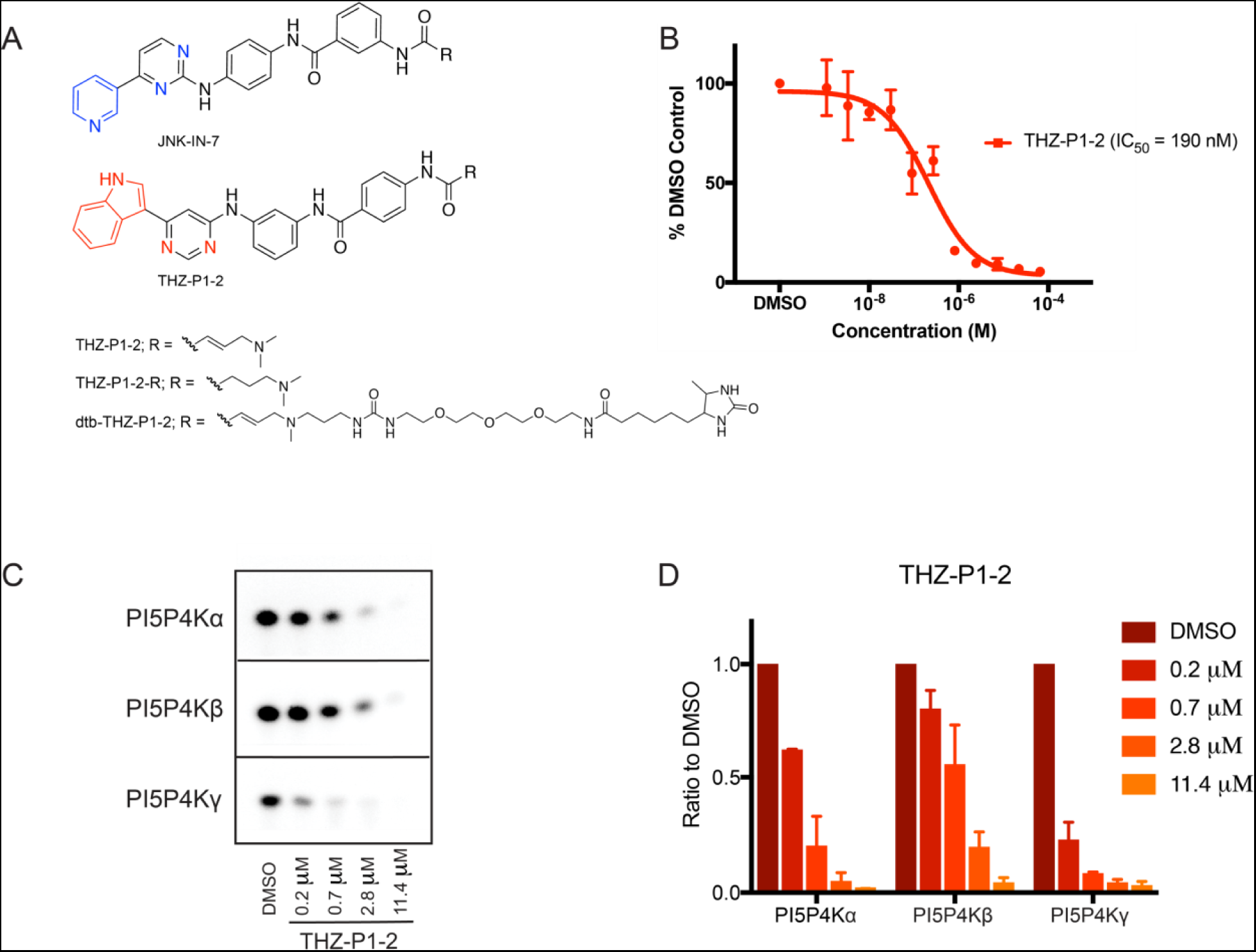
Chemoproteomic profiling and synthetic chemistry approaches reveal PI5P4K inhibitor THZ-P1-2. (A) Chemical structures of compound JNK-IN-7 as previously reported in Zhang et al (2012) and novel compound THZ-P1-2 with its reversible and desthiobiotinylated counterparts. (B) THZ-P1-2 potently inhibits PI5P4Kα kinase activity in the ADP-Glo luminescence assay. The curve obtained is representative of two independent experiments and the IC50 value is an average of values obtained from both experiments with three technical replicates each. (C) THZ-P1-2 shows potent inhibition of kinase activity of all three PI5P4K isoforms in a radiometric TLC assay measuring radiolabeled PI-4,5-P_2_. TLC image shown is representative of two independent experiments. (D) Quantification of (C). Radiolabeled PI-4,5-P_2_ spots were imaged by autoradiography and quantified by densitometry.

### THZ-P1-2 Binds in the Active Site and Covalently Modifies Cysteine Residues of the PI5P4K Kinases

Based on the initial evidence of ATP-competitive inhibition and the covalent binding of JNK-IN-7 to PI5P4Kγ, we sought to confirm covalent binding of THZ-P1-2 to each PI5P4K isoform. Upon incubating THZ-P1-2 with recombinant PI5P4Kα/β/γ protein and analyzing each sample by electrospray mass spectrometry, we discovered that the compound covalently labeled each isoform, as evidenced by the mass shift corresponding to the labeled protein, more efficiently than observed with JNK-IN-7 (Fig. 2A). Notably, THZ-P1-2 achieved 100% labeling of PI5P4Kα/β within 2 h, and PI5P4Kγ in 30 min, corroborating earlier observations in the TLC assay. This observed labeling is dependent on the compound possessing the cysteine-reactive acrylamide moiety, and we confirmed this using a non-cysteine-reactive analog of THZ-P1-2, containing a 4-(dimethylamino)butanamide warhead (Fig.1A). The reversible counterpart, denoted by THZ-P1-2-R, did not label the recombinant PI5P4K protein (Fig. S2B).

**Figure 2.**
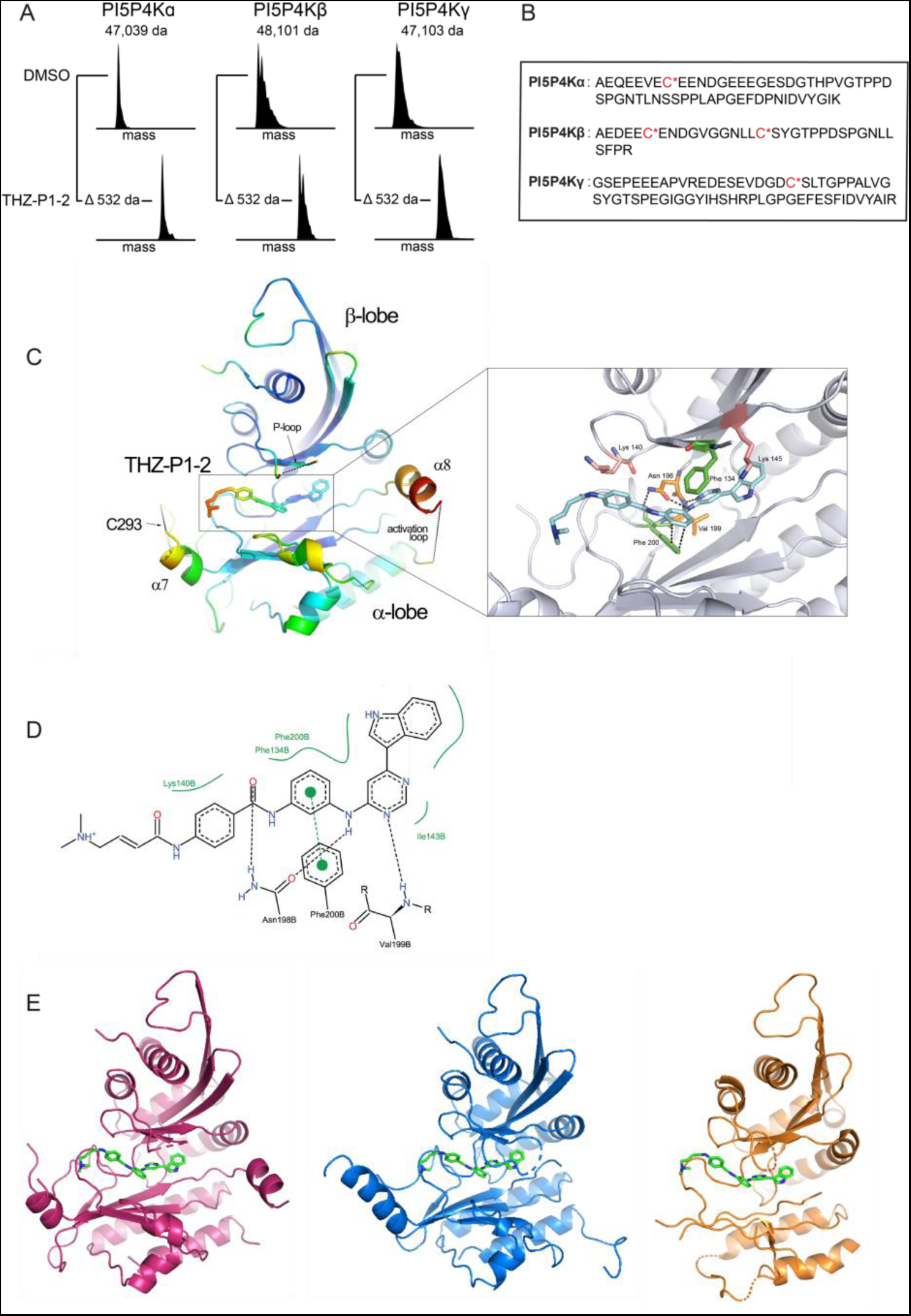
THZ-P1-2 binds covalently to all isoforms of the PI5P4K family on unique cysteine residues located on a disordered loop outside the kinase domain. (A) Electrospray mass spectrometry of recombinant PI5P4Kα/β/γ incubated with THZ-P1-2 demonstrates covalent labeling of PI5P4K isoforms. (B) Subsequent protease digestion and tandem mass spectrometry confirms that THZ-P1-2 covalently labels cysteine residues. (C) Crystal structure of PI5P4Kα in complex with THZ-P1-2, colored according to B factor and shown with covalent warhead extended out towards the covalently-targeted cysteine, C293 (labeled; not resolved in crystal structure). (D) Ligand interaction map of THZ-P1-2 with residues in the ATP-binding pocket of PI5P4Kα. (E) Modeled THZ-P1-2 binding showed for all three isoforms based on alignment of published PI5P4Kβ and γ structures with obtained PI5P4Kα structure. PI5P4Kα (magenta), PI5P4Kβ (blue), PI5P4Kγ (orange).

To ascertain the specific residues covalently modified by THZ-P1-2, we performed protease digestion and tandem mass spectrometry. Here, we identified peptides containing previously unannotated cysteine residues on a conserved disordered loop residing outside of the PI5P4K isoforms kinase ATP-binding sites – Cys293 on PI5P4Kα, Cys307 and Cys318 on PI5P4Kβ, and Cys313 on PI5P4Kγ – that were covalently modified by THZ-P1-2 (Fig. 2B, Fig. S2A). We hypothesize that this interaction is possible as the disordered loop region is brought into close proximity of the ATP-binding pocket through tertiary structure. These observations add PI5P4K to the list of kinases that possess targetable cysteines distant from the ATP-site in sequence yet accessible proximal to the ATP-site due to the overall kinase conformation (Zhang et al., 2016; Kwiatkowski et al., 2014; Browne et al., 2018).

### Elucidation of the crystal structure of THZ-P1-2 bound to PI5P4Kα

To elucidate the detailed molecular interactions involved with the covalent and non-covalent interactions of THZ-P1-2 with the PI5P4Ks, we determined a crystal structure of PI5P4Kα in complex with THZ-P1-2 to 2.21 Å resolution (Fig. 2C, Fig. S2C). The crystal structure showed crucial non-covalent interactions within the ATP-binding pocket. Asn198 forms hydrogen-bonding interactions between the carbonyl of the benzamide and the amine linker connecting the phenylenediamine with the pyrimidine moiety of THZ-P1-2. Furthermore, aromatic π-stacking interactions between the phenylenediamine and Phe200, as well as hydrophobic interactions between the indole head group and linker regions of THZ-P1-2 and several residues in the active site were detected (Fig. 2C-D). Unfortunately, as has been observed in previously published crystal structures, there was insufficient electron density to resolve the disordered loop containing the reactive Cys293 (Fig. S2D). However, it is clear from the co-structure that the cysteine-containing loop could easily traverse the region adjacent to where the acrylamide warhead of THZ-P1-2 is situated. Alignment of crystal structures of PI5P4Kβ (1BO1) and γ (2GK9) deposited in the Protein Data Bank to our resolved co-crystal structure in PyMOL allowed for a more direct comparison between the three isoforms, showing that THZ-P1-2 likely binds to the other two isoforms in a similar manner given the high degree of homology in the active sites and cysteine-containing loops (Fig. 2E).

### Kinase selectivity and cellular target engagement by THZ-P1-2

To confirm that the covalent binding observed *in vitro* is also observed in a cellular context, we synthesized a desthiobiotinylated derivative, dtb-THZ-P1-2, which maintained the biochemical potency of the parental compound (Fig. S3A). A streptavidin pulldown assay in HEK293T cell lysate confirmed that dtb-THZ-P1-2 could capture PI5P4Kα/β/γ at a concentration of 1 μM. We further demonstrated cell permeability and cellular on-target engagement of THZ-P1-2 by performing the streptavidin pulldown assay in a competitive fashion, first pre-treating HEK293T cells with the parent compound, followed by pulldown in cell lysate with the desthiobiotinylated derivative. THZ-P1-2 demonstrated the ability to compete and block pulldown of PI5P4K with dtb-THZ-P1-2 in a dose-dependent manner (Fig. 3B). We observed that engagement measured by the streptavidin pulldown assay also determined covalent binding, as pre-treatment with THZ-P1-2-R was not able to block pulldown by dtb-THZ-P1-2 (Fig. 3B). We validated THZ-P1-2’s continuous on-target engagement typical of covalent inhibitors by performing an additional pulldown with a washout at 2 hours and 4 hours, observing engagement as early as 2 hours with 1 μM compound treatment (Fig. S3B).

**Figure 3.**
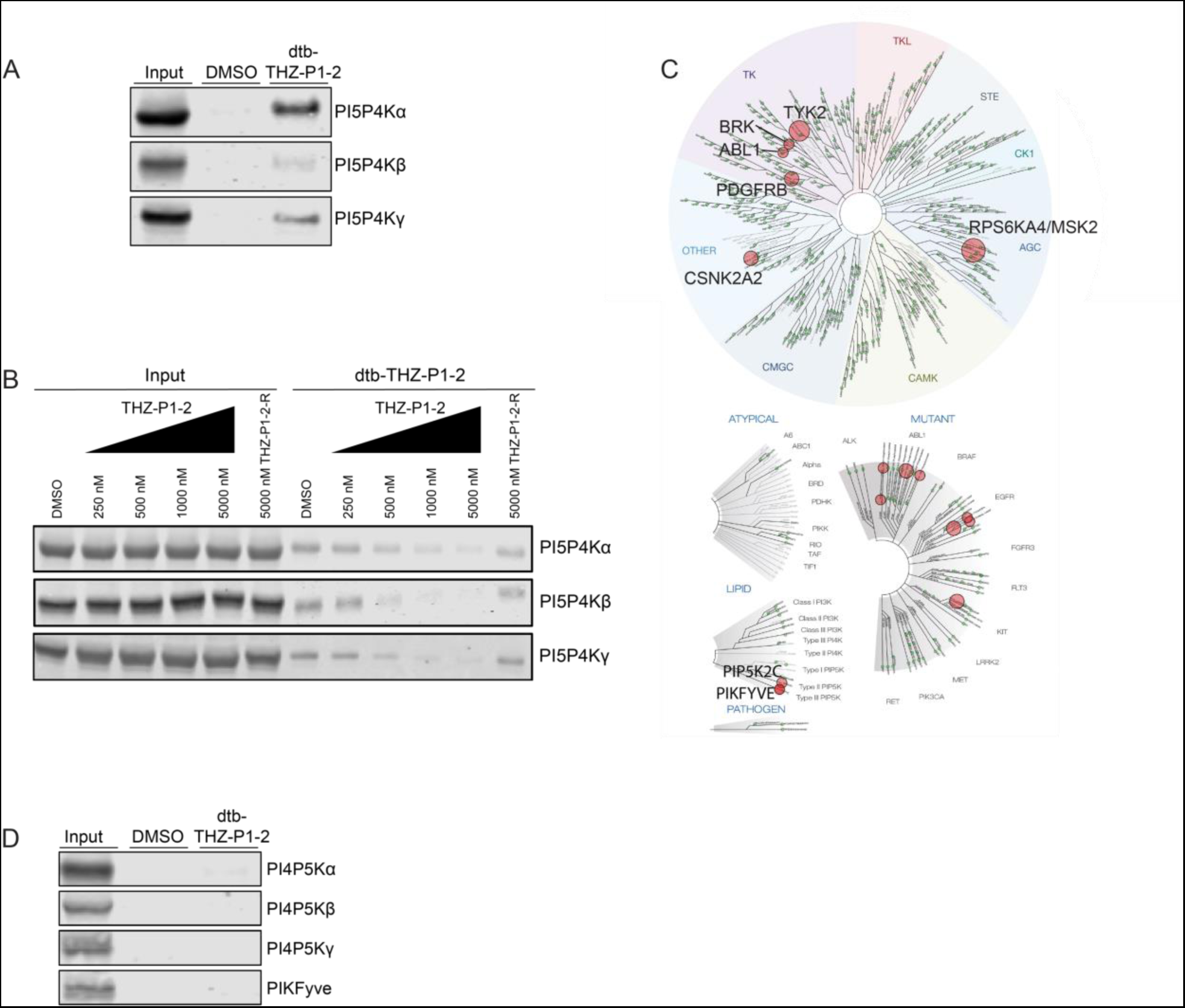
Cellular On-target Engagement and Selectivity Profile of THZ-P1-2. (A) A desthiobiotinylated analog of THZ-P1-2 irreversibly engages all PI5P4K isoforms in a streptavidin pulldown in HEK293T lysate. (B) THZ-P1-2, but not its reversible analog, exhibits dose-dependent on-target engagement of PI5P4K isoforms in a competitive streptavidin pulldown with 1 μM desthiobiotinylated-THZ-P1-2. (C) Kinome selectivity profile of THZ-P1-2. Complete table of targets included as Supplemental Data. (D) THZ-P1-2 does not engage the PI4P5Ks or PIKfyve in a streptavidin pulldown.

To broadly evaluate the kinome-wide selectivity of THZ-P1-2 we utilized the commercially available DiscoverX KINOMEScan profiling platform which revealed appreciable selectivity across the kinome with a selectivity score at a 10% DMSO control cutoff, S_(10)_, of 0.02 (indicating THZ-P1-2 bound to 2% of queried kinases at the specified cutoff) (Fig. 3C) (Karaman et al., 2008, Davis et al., 2011). We found the related lipid kinase PIKfyve to be an off-target (6.4% DMSO control). Since PIKfyve and PI4P5Kα/β/γ could reasonably be off-targets due to overall similar lipid kinase structure, we first tested THZ-P1-2 inhibitory activity on these kinases. We observed low micromolar IC_50_s against PI4P5Kα/β/γ and an IC_50_ of 40 nM on PIKfyve in a fixed time-point ADP-Glo assay (Carna) (Fig. S3C). To further assess if THZ-P1-2 biochemical inhibition of these closely related lipid kinases was maintained in cells, we evaluated dtb-THZ-P1-2 binding of these targets by streptavidin pulldown. We observed no engagement of PI4P5Kα/β/γ and PIKfyve, establishing cellular selectivity of THZ-P1-2 against these related Type 1 and Type 3 lipid kinases (Fig. 3D). Because of THZ-P1-2’s high potency on PIKfyve in the biochemical context, we took a step further to investigate its cellular activity on PIKfyve in a vacuolar enlargement assay. THZ-P1-2 produced the established PIKfyve-inhibitory phenotype in Vero cells only at a high concentration of 10 μM, 1000 x that of positive control apilimod (Cai et al., 2013; Choy & Saffi et al., 2018; Sharma & Guardia et al., 2018) which causes substantial vacuolar enlargement at 10 nM (Fig. S3D), but failed to show inhibition of PIKfyve at 1 μM. THZ-P1-2 was also unable to impair the ability of PIKfyve inhibitor apilimod to induce vacuolar enlargement when cells were either pre-incubated with THZ-P1-2 followed by apilimod, or co-incubated with both compounds (Fig. S3E).

The additional potential off-targets identified through KINOMEScan were further evaluated using the Adapta binding assay (Invitrogen). All top off-targets except for BRK and ABL1 were found to have IC_50_s in the micromolar range (Fig. S4A). Engagement of both BRK and ABL1, both of which lack cysteines in or near the vicinity of the ATP-site, were assessed in the cellular pulldown assay with dtb-THZ-P1-2 and found to have little to no pulldown, suggesting that the affinity observed in the Adapta assay may have been due to tight noncovalent binding which was not perpetuated in the cellular context (Fig. S4B). THZ-P1-2 was markedly less potent than the ABL inhibitors imatinib, nilotinib and dasatinib, at killing BCR-ABL translocation containing cell lines (K562 and KU812F) indicating that THZ-P1-2 does not effectively inhibit BCR-ABL in cells (Fig. S4C). Interestingly, even with retention of the indole head group of THZ1 and THZ531, the switch in 2,4- to 4,6-pyrimidine from JNK-IN-7 to THZ-P1-2 was sufficient to alleviate engagement of kinases targeted by related phenylaminopyrimidine acrylamides including JNK, IRAK1, PKN3, CDK7 and CDK12 when probed in a streptavidin pulldown with their respective biotinylated or desthiobiotinylated compounds (Zhang et al., 2012; Kwiatkowski et al., 2014; Browne et al., 2018) (Fig. S4D).

### THZ-P1-2 as a probe for potential PI5P4K dependencies in cancer

*PIP4K2A* was found to be essential for cell survival in AML (Jude et al., 2015) and several variant SNPs in the *PIP4K2A* locus have been associated with ALL susceptibility and leukemogenesis (Rosales-Rodríguez, et al., 2016; Urayama et al., 2018). We evaluated the sensitivity of a small panel of leukemia cell lines to THZ-P1-2, using a 72 h Cell-Titer Glo assay. We used THZ-P1-2-R as a negative compound to control for effects of THZ-P1-2 that are mediated solely by reversible binding. THZ-P1-2 demonstrated modest anti-proliferative activity in all six AML/ALL cell lines with IC_50_s in the low micromolar range. We observed approximately 10-fold IC_50_ shifts between THZ-P1-2 to THZ-P1-2-R (Fig. 4A), indicating that the presence of the PI5P4K cysteine-targeting acrylamide partially contributes to the anti-proliferative activity. ABL1 fusion genes are also heavily involved in hematological malignancies (Braekeleer et al., 2011), but the weak engagement of ABL1 in the cellular context (Fig. S4B-C), together with the covalent dependency, gave confidence that the anti-proliferative activity observed is not solely due to ABL1 inhibition.

**Figure 4.**
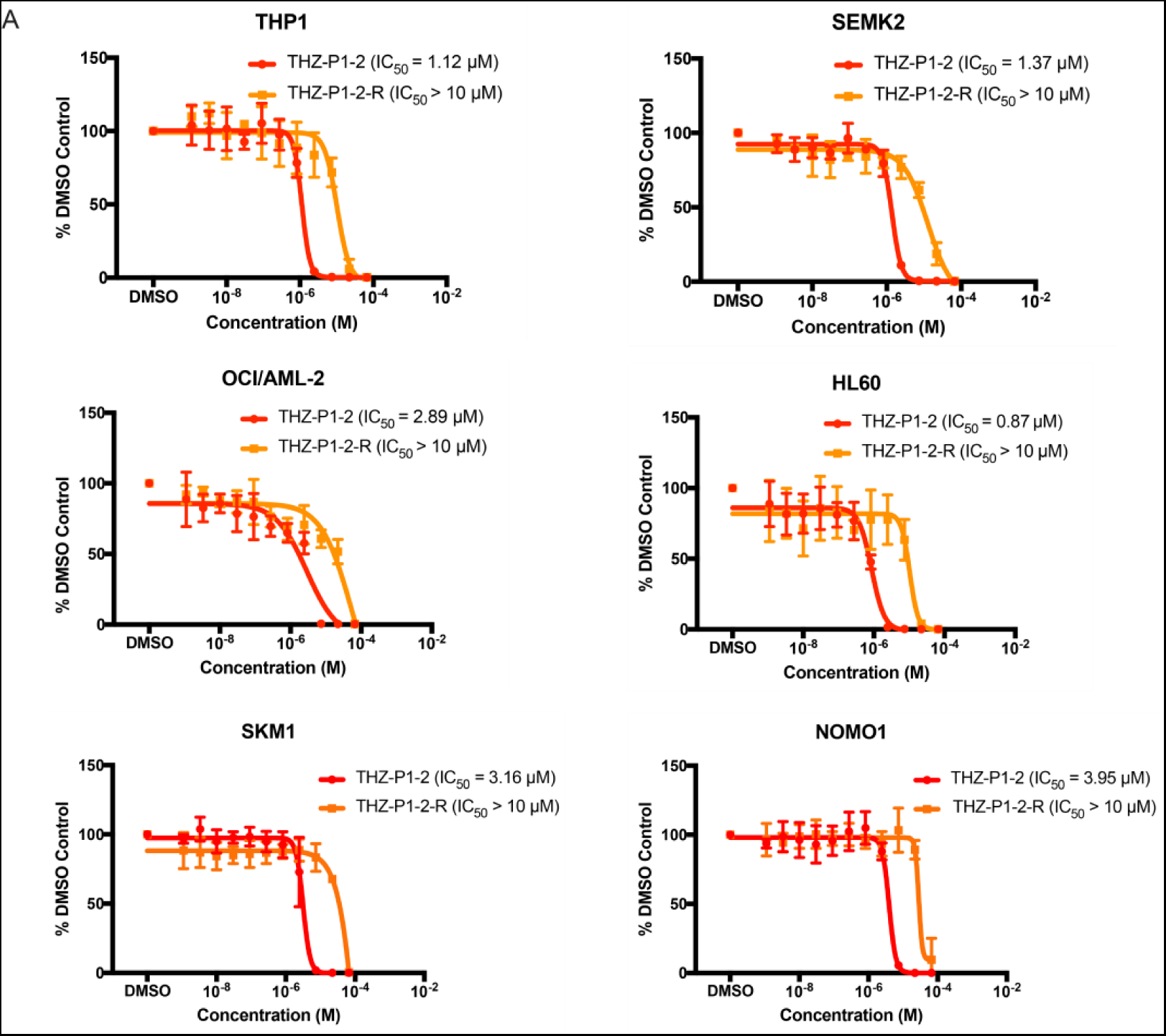
Preliminary cancer cell line profiling with THZ-P1-2 to identify potential PI5P4K dependencies. (A) A panel of AML and ALL cell lines were treated with THZ-P1-2 and THZ-P1-2-R for 72h and cell proliferation was measured with a Cell-Titer Glo luminescence assay.

### Inhibition of PI5P4K by THZ-P1-2 results in autophagy defects

Recently, PI5P4K was demonstrated to be necessary for the proper completion of autophagy in mouse models (Lundquist et al., 2018). Loss of *PIP4K2A* and *PIP4K2B* in mouse embryonic fibroblasts (MEFs) and mouse liver leads to accumulation of lysosomes and autophagosomes. Lysosomes are enlarged, fused to multiple autophagosomes, and clustered at the nuclear membrane, indicating a defect in the autophagy process. The nuclear localization of the master autophagy transcription factor TFEB was also increased and downstream transcriptional targets were upregulated, indicating an activation of the autophagy gene program (Settembre et al., 2011). However, it has yet to be demonstrated whether these defects were caused by the loss of the full-length protein or the enzymatic activity.

To validate that PI5P4K enzymatic activity is necessary for completion of autophagy, we sought to demonstrate that THZ-P1-2 phenocopies the genetic loss of PI5P4Kα and PI5P4Kβ. We first treated HeLa cells with either DMSO or THZ-P1-2 for 18 hours and stained for the lysosomal and autophagosomal markers LAMP1 and LC3B, respectively. Similar to the genetic knockouts of PI5P4K, treatment with THZ-P1-2 at concentrations as low as 250 nM results in the formation of numerous, enlarged LAMP1 puncta with fusion defects to LC3B-stained autophagosomes (Fig. 5A). This upregulation in autophagosomal/autolysosomal puncta was accompanied by a slight increase in LAMP1 protein levels with 1 μM THZ-P1-2 treatment for 24 hours, and an increase in LC3B-II in serum starvation conditions (HeLa cells cultured in media supplemented with 0.3% FBS) (Fig. S4E). We also observed an increase in nuclear TFEB accumulation (Fig. 5B-C) and the upregulation of downstream TFEB targets with THZ-P1-2 treatment (Fig. 5F).

**Figure 5.**
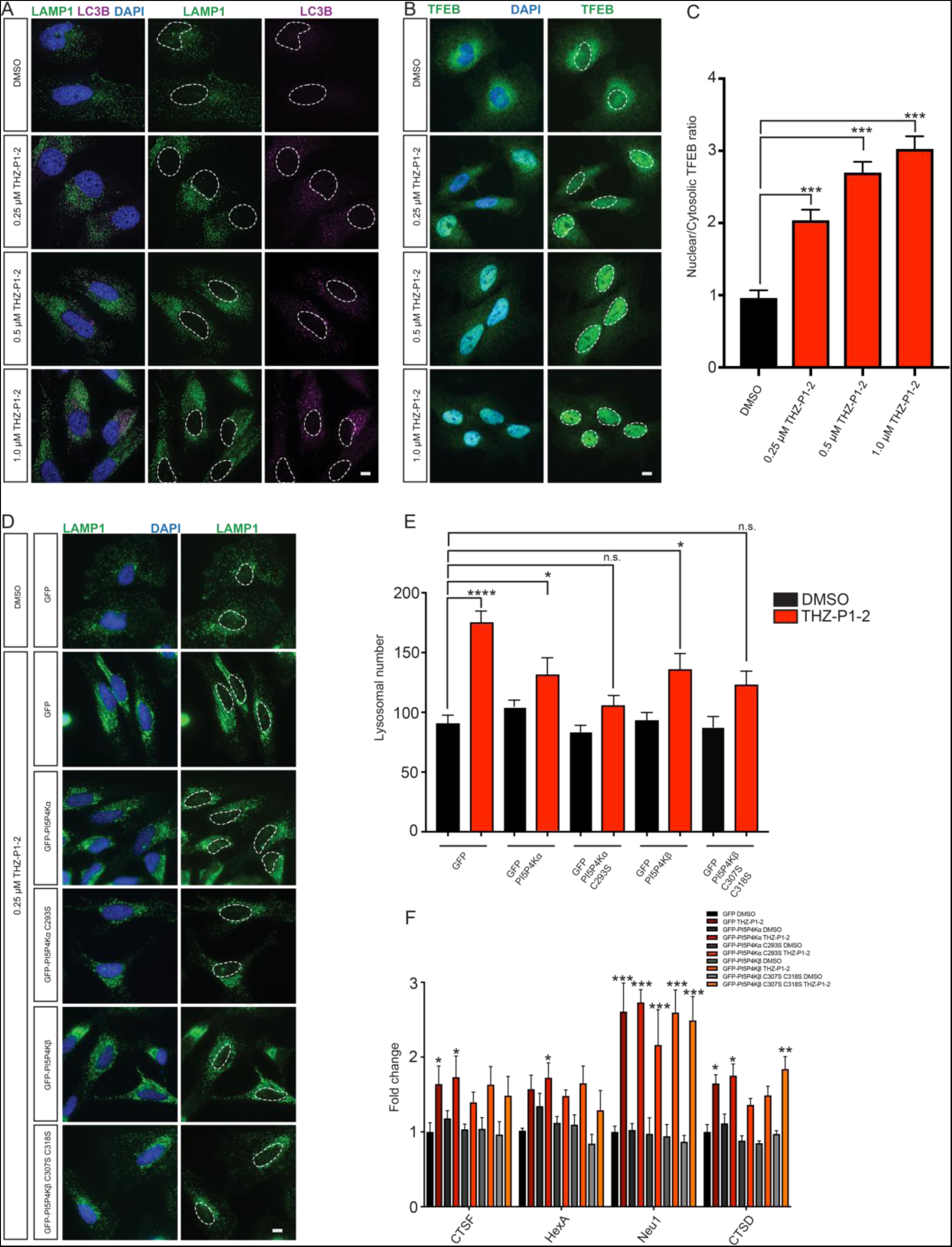
Inhibition of PI5P4K with THZ-P1-2 leads to lysosomal-autophagosomal defects and increased TFEB activation. (A) Similar to genetic loss of *PIP4K2A/B*, inhibition of PI5P4K activity with THZ-P1-2 increases LAMP1-positive lysosomal size, number and contact with LC3B-positive autophagosomes. HeLa cells were cultured overnight with either DMSO, 0.25 μM, 0.5 μM or 1.0 μM THZ-P1-2 and stained for LC3B (magenta) or LAMP1 (green) with nuclei in blue. Scale bars, 10 μM. (B) Inhibition of PI5P4K activity with THZ-P1-2 increases TFEB nuclear localization. HeLa cells were cultured overnight with either DMSO, 0.25 μM, 0.5 μM or 1.0 μM THZ-P1-2 and stained for TFEB (green) with nuclei in blue. Scale bars, 10 μM. (C) Quantification of results in (B). The intensity of TFEB immunofluorescence was quantified in the nucleus and the cytoplasm, and used to calculate the ratio. Statistical significance determined by ANOVA (***p < 0.0005) with Dunnett multiple comparison post-test. Each group was compared to control control HeLa cells treated with DMSO, (n ≥ 30). (D) Expression of PI5P4K cysteine to serine mutants alleviates lysosomal dysfunction induced by THZ-P1-2 treatment. HeLa cells were infected with viruses expressing GFP, GFP-PI5P4Kα, GFP-PI5P4Kα C293S, GFP-PI5P4Kβ, or GFP-PI5P4Kβ C307S C318S and treated with either DMSO or 250 nM THZ-P1-2 overnight. Expression of both the PI5P4Kα and PI5P4Kβ cysteine to serine mutants alleviated dysfunctional lysosomes (green) with nuclei in blue. Scale bars, 10 μM. (E) Quantification of results in (D). The number of lysosomes was quantified per cell. Statistical significance determined by ANOVA (***p < 0.0005) with Dunnett multiple comparison post-test. Each group was compared to control HeLa cells expressing GFP and treated with DMSO, (n ≥ 30). (F) HeLa cells expressing GFP, GFP-PI5P4Kα, GFP-PI5P4Kα C293S, GFP-PI5P4Kβ, or GFP-PI5P4Kβ C307S C318S were treated with either DMSO or 250 nM THZ-P1-2 overnight for 16 hours and subsequently harvested for qPCR of TFEB targets. Fold change is calculated by comparison to HeLa cells expressing GFP treated with DMSO. *p < 0.05, Student’s t-test, (n ≥ 8).

We next investigated whether the effects of THZ-P1-2 are specific to PI5P4K by rescuing the autophagy defects with the expression of PI5P4Kα/β Cys-Ser mutant isoforms, which are unable to bind the inhibitor covalently. We infected HeLa cell lines to stably express a GFP-only vector, GFP-tagged PI5P4Kα/β, and GFP-tagged PI5P4K Cys-Ser mutants (α: C293S; β: C307,318S;). We treated the cells for 18 hours with either DMSO or THZ-P1-2. As observed with uninfected cells, cell lines expressing GFP alone saw an increase in lysosome/autophagosome dysfunction with THZ-P1-2 treatment (Fig. 5D-E). These effects were partially ameliorated by expression of the PI5P4K constructs and even more so by the cysteine to serine mutants (Fig. 5D-E). The PI5P4K Cys-Ser mutants also partially rescued the nuclear localization of TFEB and the expression of downstream TFEB target genes (Fig. 5D-F).

## Discussion

The PI5P4Ks comprise a family of lipid kinases that regulate levels of their substrate and product PI-5-P and PI-4,5-P_2_, maintaining compartment-specific concentrations of these phosphoinositides at a subcellular level. Initially thought to be mainly a PI-5-P modulator within a noncanonical PI-4,5-P_2_ generation pathway, the PI5P4Ks have garnered a lot of interest in recent years because of their purported importance in specific disease contexts, spurring efforts to develop PI5P4K inhibitors. In this study, we designed and characterized THZ-P1-2, a first-in-class covalent inhibitor of the PI5P4K family. THZ-P1-2, a small molecule containing an electrophilic acrylamide moiety capable of undergoing a Michael addition, bound in the active site and irreversibly inhibited the enzymatic activity of these kinases. A covalent targeting strategy with THZ-P1-2 resulted in prolonged on-target engagement in cells and modest anti-proliferative activity in a panel of AML and ALL cell lines, while retaining a reasonable selectivity across the kinome. Lastly, we found that THZ-P1-2 treatment phenocopied the effects of genetic deletion of PI5P4Kα/β/γ in mice by causing autophagy defects in HeLa cells.

Our initial point of compound development was JNK-IN-7, a previously reported covalent JNK inhibitor that was found to be capable of binding to PI5P4Kγ by KiNativ profiling. Originally inspired by the imatinib core structure, this “privileged” kinase scaffold served as a promising starting point for developing a selection of analogs with chemical modifications with which to explore structure-activity relationships. Among this panel of molecules (to be published at a later date), we found a critical feature resulting in superior potency to be a switch from a 2,4-pyrimidine to a 4,6-pyrimidine hinge-interacting motif. We established THZ-P1-2’s inhibitory activity with an apparent IC_50_ of 190 nM on PI5P4Kα in a fixed time-point assay, the isoform with the highest kinase activity and thus easiest to analyze using an ATPase assay (ADP-Glo). We confirmed that THZ-P1-2 specifically prevents phosphate transfer to the PI-5-P substrate at nanomolar concentrations for all isoforms with an orthogonal TLC assay measuring radiolabeled PI-4,5-P_2_ formation.

THZ-P1-2 covalently labels all three PI5P4K isoforms, with the reactive cysteine residues located on analogous disordered loops of unknown function outside the kinase domain. There has been a resurgence of interest in covalent inhibitors, especially with recent clinical successes of cysteine-directed covalent BTK inhibitor ibrutinib (Pan et al., 2007), EGFR inhibitors neratinib (Rabindran et al., 2004), afatinib (Li et al., 2008), and osimertinib (Cross et al., 2014). Covalent irreversible inhibitors characteristically exhibit an enhanced potency and prolonged pharmacodynamics due to permanent disabling of kinase enzymatic activity, and high selectivity as covalent inhibitors will only bind irreversibly to a kinase with an appropriately placed cysteine, within the trajectory of the electrophilic warhead (Liu et al., 2013). Our approach in developing probes THZ1 against CDK7 (Kwiatkowski et al., 2014), TH531 against CDK12/13 (Zhang et al., 2016) and most recently JZ128 against PKN3 (Browne et al., 2018) showcased to great effect the modification of a reactive cysteine residue outside the active site, widening the scope of “accessible” cysteines. We sought to apply these principles and strategies towards PI5P4K. Our study validates the PI5P4Ks as additional candidate kinases with cysteines that are distant from the active site in sequence but brought close to the ATP-binding site due to kinase tertiary structure, rendering them amenable to covalent targeting. Moreover, our approach in targeting these unique cysteines provides a proof-of-concept in using a combination of inhibitor covalent and noncovalent affinities to achieve selectivity, due to a lack of equivalent off-target cysteines across the kinome.

Structural insights into the binding mode of THZ-P1-2 with PI5P4Kα were obtained through a co-crystal structure. Besides the noncovalent interactions highlighted in the results, hydrogen-bonding between the backbone of Val199 and the 4,6-pyrimidine may account for the increased potency compared to analogs with a 2,4-pyrimidine, especially if they contain substituents on the heterocycle linker. Notably, the Phe205 residue that engages in aromatic π-stacking interactions with the phenylenediamine moiety of THZ-P1-2 is conserved among the PI5P4Ks and has been identified as the residue responsible for the GTP-sensing ability of PI5P4Kβ (Sumita et al., 2016). This co-crystal structure also serves as an important resource for the scientific community, enabling structure-guided design of PI5P4K inhibitors with improved properties. Cell permeability and dose-dependent on-target engagement of THZ-P1-2 was validated in HEK293T cells, chosen for their robust expression of all three PI5P4K isoforms, and the necessity for covalency verified through observing effects after washout and in anti-proliferation assays alongside THZ-P1-2-R.

Our biochemical selectivity profiling of THZ-P1-2 indicates that though the compound exhibits satisfactory selectivity, there are some off-targets observed. The most potent off-targets, such as ABL1, PIKfyve and BRK, were not engaged by dtb-THZ-P1-2 in a cellular pull-down assay. PIKfyve was inhibited by THZ-P1-2 only at concentrations of 10 μM or greater, higher than concentrations relevant for observable PI5P4K targeting. These results indicate that THZ-P1-2 does have some off-target activity on PIKfyve, but maintains a preference for the PI5P4Ks, again possibly due to a combination of covalent and non-covalent binding modes.

A principal part of our study was to further confirm novel functions of the PI5P4Ks beyond their asserted primary role of non-canonical PI-4,5-P_2_ generation. Treatment with THZ-P1-2 phenocopied genetic deletion of PI5P4K in causing autophagosome and lysosome disruption, TFEB nuclear localization and TFEB target gene upregulation, indicating that PI5P4K kinase activity is required for the role of these kinases in autophagy. Cancer cell line profiling demonstrates AML/ALL cancer cell lines to be sensitive to THZ-P1-2 covalent targeting. Consistent with findings that PI5P4K mediates autophagy in times of nutrient stress (Lundquist et al., 2018) our studies using THZ-P1-2 as a tool suggest that the PI5P4Ks display an induced essentiality by facilitating autophagy as an energy source in a metabolically-stressed environment such as cancer (Kimmelman & White, 2017). Additional studies are required to fully dissect the connection between PI5P4K inhibition, autophagy disruption and subsequent effects on cell viability, as well as investigate the sensitivities of various cancer types to specific PI5P4K inhibition. THZ-P1-2 may be a useful probe of the pharmacological consequences of covalent inhibition of PI5P4K that can complement genetic approaches (such as CRISPR- or RNAi-based screening) to identify unique vulnerabilities in cancer.

In conclusion, we have characterized a novel potent and selective covalent PI5P4K inhibitor that exhibits durable cellular pharmacodynamics, disruption of autophagy, and modest anti-proliferative activity against leukemia-derived cell lines. THZ-P1-2, THZ-P1-R and the inhibitor resistant PI5P4K Cys to Ser mutants described in this manuscript provide a chemical biology toolbox serving as a valuable resource with which to investigate the pharmacological consequences of covalent inhibition of PI5P4Ks. Our discovery of THZ-P1-2 may be used as a model for targeting other understudied gene families, especially those bearing targetable cysteines in unique sites. Lastly, our small-molecule strategy for disrupting autophagy through covalent PI5P4K inhibition exposes a new Achilles heel and indicates that exploiting this metabolic vulnerability may be a viable therapeutic strategy in cancer and potentially other autophagy-addicted disorders.

### Significance

Our discovery and characterization of a novel pan-PI5P4K inhibitor, THZ-P1-2, that covalently targets cysteines on a disordered loop in PI5P4Kα/β/γ presents a new way to target kinases, especially noncanonical lipid kinases, by taking advantage of unique unannotated cysteines outside the kinase domain. THZ-P1-2 demonstrates cellular on-target engagement with limited off-targets across the kinome, displaying requisite potency and selectivity to be a useful tool compound. The sensitivity of leukemia cancer cells to THZ-P1-2 covalent inhibition is evidence of the potential of THZ-P1-2 to be used more universally in larger scale cancer cell line screens to identify PI5P4K dependencies in other cancer types, or other diseases beyond cancer. This can be done in parallel with typical genetic approaches (such as CRISPR- or RNAi-based screening) to identify unique vulnerabilities in a cheaper, faster, less labor-intensive manner. One of the major advantages of THZ-P1-2, as with covalent inhibitors in general, is the ability to verify on-target activity through inhibitor-resistant cysteine-to-serine mutations. In the case of the PI5P4K family of enzymes which has three isoforms, the utility of having both covalent and reversible compounds is extremely valuable, where both compounds can be used in combination to broadly assess PI5P4K dependencies in disease, with follow-up mutant rescue studies to verify the PI5P4K contribution once the scope is narrowed down. Finally, THZ-P1-2-induced autophagy disruption phenocopying the effects of PI5P4K genetic deletion lends confidence to the on-target activity of the compound and presents a novel way to target autophagy-lysosome homeostasis in disease. Our studies demonstrate that PI5P4Ks are tractable targets, with THZ-P1-2, its reversible and desthiobiotinylated analogs, and wild-type and mutant cell lines as a useful toolbox to further interrogate the therapeutic potential of PI5P4K inhibition, providing a new starting point for drug discovery campaigns for these lipid kinases in cancer metabolism and other autophagy-dependent disorders.

## Supporting information

Supplemental Data 1. DiscoverX kinome selectivity list for THZ-P1-2

## Acknowledgements

The authors acknowledge generous support from NIH grants 1U19AI109740 (to J.M.C.), R21 CA178860 (to J.A.M.), R21CA188881 (to J.A.M.), R35 CA197588 (to L.C.C.), R01 GM041890 (to L.C.C.), U54 U54CA210184 (to L.C.C.), R01 CA197329 (to N.S.G. and S.D.P.), the Dana-Farber Strategic Research Initiative (to J.A.M.) and the Breast Cancer Research Foundation (to L.C.C.). Synchrotron data collection was based upon research conducted at the Advanced Photon Source on the Northeastern Collaborative Access Team beamlines (NIGMS P41 GM103403).

## Author contributions

L.C.C., H.S., T.H.Z. and N.S.G. conceived original project. S.C.S. designed and performed the majority of experiments. T.H.Z., B.S.J., M.F.H. and T.D.M. designed and synthesized compounds. C.M.B. and S.B.F. conducted MS experiments. M.R.L. performed autophagy experiments. H.S.S. performed protein purification and crystallography experiments. H.S.S. and S.D.P. performed crystallography data collection and analysis. M.N.P., P.K., D.G.W., and T.J.Y. performed experiments. N.P.K. provided expertise and feedback. J.M.C., J.A.M., S.D.P., L.C.C. and N.S.G. supervised the research. L.C.C. and N.S.G. secured funding. S.C.S. wrote the manuscript, with guidance from F.M.F., T.H.Z. and N.P.K.. S.C.S., H.S., F.M.F., T.D.M., and N.S.G. revised the manuscript. All authors read and provided feedback on the manuscript.

## Materials & Methods

### Chemistry

All solvents and reagents were used as obtained. ^1^H-NMR spectra were recorded with a Varian Inova 600 NMR spectrometer and referenced to DMSO. Chemical shifts are expressed in ppm. Mass spectra were measured with Waters Micromass ZQ using an ESI source coupled to a Waters 2525 HPLC system operating in reverse mode with a Waters Sunfire C18 5 μm, 4.6 × 50 mm column. Purification of compounds was performed with either a Teledyne ISCO CombiFlash Rf system or a Waters Micromass ZQ preparative system. The purity was analyzed on a Waters LC-MS Symmetry (C18 column, 4.6 × 50 mm, 5 μM) using a gradient of 5%–95% methanol in water containing 0.05% trifluoroacetic acid (TFA). Detailed synthetic schemes and characterization data below (see Scheme 1).

### Cell Culture

All cell culture was performed using standard techniques. HEK293T and HeLa cells were cultured in Dulbecco’s Modified Eagle’s medium (DMEM) supplemented with 10% FBS and 1% penicillin/streptomycin (Gibco). Leukemia cells were cultured according to ATCC or DSMZ recommendations. All cells were cultured at 37°C and 5% CO_2_.

### ADP-Glo kinase assays

ADP-Glo assay protocol was modified from Davis et al. (2013). DPPS and PI5P (Echelon Biosciences) were dissolved in DMSO (333 μL/mg) and mixed by sonication and vortexing in a ratio of 2:1 DPPS:P15P. First, 63 µL of DMSO was added to 1255 µL of buffer 1 (30 mM Hepes pH 7.4, 1 mM EGTA, 0.1% CHAPS) and 2868 µL of buffer 2 (46 mM Hepes pH 7.4, 0.1% CHAPS). Volumes were multiplied according to number of assays to be run. PI5P4Kα enzyme was added to the buffer mixture at 32 nM concentration (predetermined on a batch-by-batch basis to give maximal signal to background). 10 µL was dispensed into white 384-well plates (Corning #3824). Then, 100 nL of compounds in DMSO were transferred by a pintool (JANUS, PerkinElmer). To initiate the reaction, 5 µL of ATP (Promega) at 15 µM (final assay concentration of 5 µM) and PI5P/DPPS (at concentrations of 0.06 µg/µL and 0.12 µg/µL respectively) in buffer 3 (20 mM Hepes pH 7.4, 60 mM MgCl_2_, 0.015 mM ATP and 0.1% CHAPS) was added to each well. The final concentration of DMSO in the reaction was less than 5%. The resulting mixture was incubated at room temperature in the dark for one hour, at which time 5 µL of ADP-Glo reagent 1 were added to stop the reaction and remove any remaining ATP. After a 45-minute incubation, 10 µL of the ADP-Glo reagent 2 were added and allowed to incubate for 30 minutes. The luminescence was then read on an EnVision 2104 Multilabel Plate Reader (PerkinElmer). IC_50_s were determined using the GraphPad Prism nonlinear regression curve fit.

### Radiometric kinase assays

The PI5P4K assay was carried out as described in Rameh et al. (1997). Briefly, the kinase reaction was carried out in a total of 70 µL of kinase buffer (50 mM HEPES pH 7.4, 10 mM MgCl_2_) with 0.1 µg of purified PI5P4K protein (preincubated with DMSO or compound at various concentrations), followed by addition of 20 µL of the resuspended lipids (sonicated and vortexed), 10 µM non-radiolabeled ATP, and 10 µCi [γ-^32^P]-ATP for 10 minutes at room extracted by adding 100 µL of methanol/chloroform (approximately 1: temperature. The reaction was terminated by adding 50 µL of 4 N HCl. Phosphoinositides were extracted by adding 100 μL of methanol/chloroform (approximately 1:1, vol:vol) mix and subjected to TLC (thin-layer chromatography) separation using heat-activated 2% oxaloacetate-coated silica gel 60 plates (20 cmx20 cm, EMD Millipore) and a 1-propanol/2 M acetic acid (65:35, vol:vol) solvent system. The radiolabeled product, PI(4,5)P_2_, was quantified with a Phosphorimager (Molecular Dynamics, STORM840, GE Healthcare).

### Immunoblotting

Cells were lysed with Pierce IP Lysis buffer supplemented with a cOmplete™ Mini Protease Inhibitor Cocktail tablet (Roche) on ice for 60 min. Lysates were clarified by centrifugation at 20,000 × *g* for 15 min at 4°C. Protein concentration was determined by a BCA assay (Pierce), and all samples were run with equal total protein content. The following antibodies were used in this study: PI5P4Kα (5527; Cell Signaling), PI5P4Kβ (9694; Cell Signaling), PI5P4Kγ (HPA058551; Sigma-Aldrich),PI4P5Kα (9693; Cell Signaling), PI4P5Kβ (12541-1-AP; Proteintech), PI4P5Kγ (3296; Cell Signaling), PIKfyve (MABS522; EMD Millipore), c-Abl (sc-56887; Santa Cruz), Brk (ab137563; Santa Cruz), IRAK1 (4504; Cell Signaling), PKN3 (NBP1-30102; Novus Biologicals), SAP/JNK (9252; Cell Signaling), CDK7 (2916; Cell Signaling), CDK12 (11973; Cell Signaling), LAMP1 (9091; Cell Signaling), LC3B (2773; Cell Signaling), tubulin (3873; Cell Signaling). Imaging was performed by detection of fluorescently labeled infrared secondary antibodies (IRDye) on the Odyssey CLx Imager (LI-COR).

### Pulldown assays

HEK293T cells were lysed with Pierce IP Lysis buffer supplemented with a cOmplete™ Mini Protease Inhibitor Cocktail tablet (Roche) on ice for 60 min. Lysates were clarified by centrifugation at 20,000 × *g* for 15 min at 4°C. Protein concentration was determined by a BCA assay (Pierce), and all samples were equalized for protein content. DMSO or 1 μM dtb-THZ-P1-2 was added and rotated overnight at 4°C. Samples were rotated for 2h at room temperature to enhance covalent binding and then incubated with streptavidin resin for 2h at 4°C. Beads were washed with lysis buffer five times, resuspended in 1x LDS sample buffer, boiled at 95°C for 10 min and subjected to immunoblotting. For competitive pulldowns, HEK293T cells were grown to 90-95% confluence, pretreated for 6h with DMSO or varying concentrations of THZ-P1-2, washed twice with cold PBS, lysed, and streptavidin pulldown conducted as described above with normalized samples.

### Mass Spectrometry

For intact MS analysis, recombinant PI5P4K protein was incubated with DMSO or 5 μM inhibitor (JNK-IN-7, THZ-P1-2 or THZ-P1-2-R) for 2 h at 37°C and analyzed by LC-ESI-MS essentially as described in Zhang et al. (2012). In each analysis, 5 μg protein was injected onto a self-packed reversed phase column (1/32” O.D. × 500 um I.D., 5 cm of POROS 10R2 resin). After desalting for four minutes, protein was eluted with an HPLC gradient (0–100% B in 4 min, A = 0.2M acetic acid in water, B = 0.2 M acetic acid in acetonitrile, flow rate = 10 μL/min) into an LTQ ion trap mass spectrometer (ThermoFisher). Mass spectra were decon-voluted using MagTran1.03b2 software. To determine the site of modification, the samples used for intact analysis were reduced with TCEP (10 mM final concentration), alkylated with iodoacetamide (22.5 mM final concentration) and digested with Asp-N or Glu-C (37 °C, overnight) and analyzed by nanoLC-MS. Digested peptides were injected onto the precolumn (4 cm POROS 10R2, Applied Biosystems) and eluted with an HPLC gradient (NanoAcquity UPLC system, Waters, Milford, MA; 10–70% B in 60 min; A = 0.1 M acetic acid in water, B = 0.1M acetic acid in acetonitrile). Peptides were resolved on a self-packed analytical column (50 cm Monitor C18, Column Engineering, Ontario, CA) and introduced to the mass spectrometer (LTQ Orbitrap XL) at a flow rate of ∼30 nL/min (ESI spray voltage = 3.2 kV). The mass spectrometer was programmed to perform data-dependent MS/MS on the five most abundant precursors (35% collision energy) in each MS1 scan (image current detection, 30K resolution, *m*/*z* 300–2000). MS/MS spectra were matched to peptide sequences using Mascot (version 2.2.1) after conversion of raw data to .mgf using multiplierz scripts. Precursor and peptide ion mass tolerances were 10 ppm and 0.6 Da, respectively. Supplemental Intact MS Spectra for representative raw MS spectra used for charge deconvolution to be included separately.

### Protein expression and purification

A construct of human PI5P4Kα covering residues 35-405 in the pNIC28Bsa4 vector (Addgene #42494) was overexpressed in E. coli BL21 (DE3) in TB medium in the presence of 50 μg/ml of kanamycin. Cells were grown at 37°C to an OD of 0.7, induced overnight at 17°C with 400 μM isopropyl-1-thio-D-galactopyranoside, collected by centrifugation, and stored at −80°C. Cell pellets were resuspended in buffer A (50 mM sodium phosphate, pH 7.4, 500 mM NaCl, 10% glycerol, 20 mM Imidazole, and 14 mM BME), lysed by sonication, and the resulting lysate was centrifuged at 16,000 xg for 30 min. ∼5mL Ni-NTA beads (Qiagen) were mixed with lysate supernatant for 45 min, washed with buffer A, and eluted with buffer B (50 mM sodium phosphate, pH 7.4, 500 mM NaCl, 10% glycerol, 300 mM Imidazole, and 14 mM BME). The eluted sample was gel-filtered through a Superdex-200 16/600 column in buffer C (20 mM HEPES, pH 7.5, 500 mM NaCl, 10% glycerol, 10mM DTT, and 1mM TCEP). Protein fractions were pooled, concentrated, and stored at −80°C.

### Crystallization

A sample of 400 μM protein and 500 μM THZ-P1-2 was co-crystallized in 20% PEG3350 and 200 mM NaMalate by sitting-drop vapor diffusion at 20°C. Crystals were transferred briefly into crystallization buffer containing 25% glycerol prior to flash-freezing in liquid nitrogen.

### Data collection and structure determination

Diffraction data from complex crystals were collected at beamline 24ID-E of the NE-CAT at the Advanced Photon Source at the Argonne National Laboratory. Data sets were integrated and scaled using XDS (Kabsch, 2010). Structures were solved by molecular replacement using the program Phaser (McCoy et al., 2007) and the search model PDB entry 2YBX. Iterative model building, ligand fitting, and refinement using Phenix (Adams et al., 2010) and Coot (Emsley & Cowtan, 2004) led to models with excellent statistics, shown in Table S1.

### Proliferation assays

Cells were plated in 96-well or 384-well format and treated with indicated compounds at varying concentrations for 72h. Anti-proliferative effects of compounds were assessed using the Cell Titer Glo assay kit (Promega) with luminescence measured on an EnVision 2104 Multilabel Plate Reader (PerkinElmer). IC_50_s were determined using the GraphPad Prism nonlinear regression curve fit.

### qRT-PCR

Total RNA was prepared using RNeasy (Qiagen). cDNA was synthesized using Superscript Vilo (Thermo) and qRT-PCR performed utilizing Fast SYBR green (Thermo) and the Realplex Mastercycler (Eppendorf). For a list of primers used see Table S1. Isolation of mRNA and qPCR was performed as follows. 200,000 cells were plated in 6-well plastic dishes. 24 hours later, the RNA in the lysates was extracted using the RNeasy protocol. The RNA was resuspended in 50 μl H_2_O at a concentration of 1 μg/μL. cDNA was transcribed using the SuperScript Vilo. The sequences of the oligonucleotides used as primers in the PCR reactions are given in Table S1. The genes that were quantified here were previously shown to be regulated by TFEB (Perera et al., 2015).

### Fluorescence microscopy

HeLa were grown on glass coverslips pre-treated with poly-d-lysine. When indicated cells were treated with 250-1000 nM THZ-P1-2 for 16 hours. Adherent cell lines were rinsed with phosphate-buffered saline, pH 7.4 (PBS) and fixed with 4% paraformaldehyde (PFA) in PBS for 15 minutes at room temperature. After fixation, the cells were permeabilized for 10 minutes with PBS/0.1% Triton X-100, blocked for 30 minutes in blocking buffer (PBS with 3% BSA) and labeled with primary antibodies in blocking buffer for 1 hour at room temperature or overnight at 4°C. Alternatively, for staining, LC3B and LAMP1 (ab25245; Abcam) cells were fixed/permeabilized in −20 MeOH for 20 minutes. Coverslips were washed three times with blocking buffer and incubated with Alexa Fluor-conjugated goat secondary antibodies in blocking buffer for 1 hour at room temperature. After incubation with secondary antibodies, coverslips were washed three times with PBS, once with water, and then mounted on a glass microscope slide with Prolong Gold with DAPI (Thermo). The following primary antibodies were used: TFEB (SAB4503154; Sigma), LC3B (3868; Cell Signaling), LAMP1 (ab25245; Abcam). Alexa Fluor-conjugated secondary antibodies (Thermo) were used at 1:1000. Fluorescent and phase contrast images were acquired on a Nikon Eclipse Ti microscope equipped with an Andor Zyla sCMOS camera. Within each experiment, exposure times were kept constant and in the linear range throughout. When using the 60x and 100x oil immersion objectives, stacks of images were taken and deconvoluted using AutoQuant (Media Cybernetics).

### Generation and transduction of GFP *PIP4K2* lentiviral constructs

Replication-deficient lentiviruses were prepared using a third-generation lentiviral system. GFP was inserted by itself or upstream of either human *PIP4K2A* or *PIP4K2B* in a lentiviral vector. Cysteine to Serine point mutations were made using the Quikchange XL II site-directed mutagenesis kit (Agilent). Virus was generated by cotransfecting three helper plasmids (pLP1, pLP2, pVSV-G) and the vector containing the gene of interest with *cis*-acting sequences for proper packaging were used to generate pseudovirions. A subconfluent culture of HEK293T cells was transfected using the CalPhos Mammalian Transfection Kit (Clontech). The titer of each virus was determined using HEK293T cells. Viral supernatant was then used to express PI5P4Ks in MEFs.

### Virus production and infection

293T packaging cell line was used for retroviral amplification. In brief, viruses were collected 48 hours after infection, filtered, and used for infecting cells in the presence of 8μg/ml polybrene (Sigma), prior to puromycin selection or upon cell sorting for GFP. The packaging plasmid used for retroviral infection was pCL-Eco (Naviaux, Costanzi, Haas, & Verma, 1996).

**Scheme 1.**
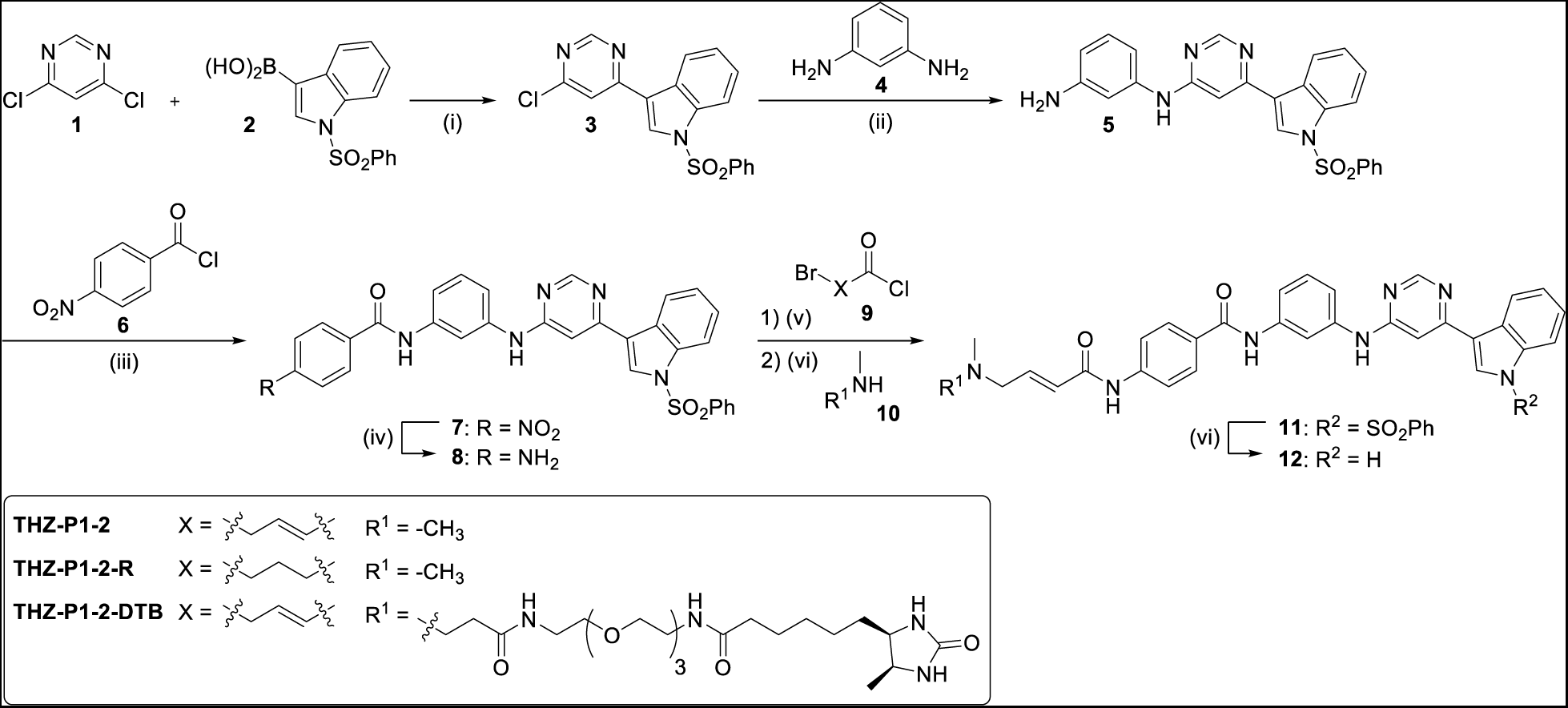
Synthetic route of THZ-P1-2/THZ-1-2-R/dtb-THZ-P1-2.^*a*^. ^*a*^ Reagents and conditions: (i) NaHCO_3_, Pd(PPh_3_)_2_Cl2, ACN/H_2_O (4:1, v/v), 90 °C, overnight, 61 % (ii) DIEA, NMP, 150 °C, overnight, 95 % (iii) pyridine, 80 °C, 3 h (iv) SnCl_2_·2H_2_O, EtOAc/MeOH (5:2, v/v), 80 °C, overnight, 69 % over 2 steps (v) 1) DIEA, ACN, 0 °C, 5 min. 2) THF, rt, 2 h (vi) 1 m NaOH/1,4-dioxane (1:1, v/v), rt, 6 h, 7-73 % over 3 steps.

THZ-P1-2: MS m/z 532.24 [M+H]^+. 1^H NMR (400 MHz, DMSO-*d*_6_) δ 11.73 (s, 1H), 10.50 (s, 1H), 10.19 (s, 1H), 9.55 (s, 1H), 8.63 (s, 1H), 8.27 (d, *J* = 7.2 Hz, 1H), 8.20 (s, 1H), 8.15 (d, *J* = 2.8 Hz, 1H), 8.00 (d, *J* = 8.7 Hz, 2H), 7.83 (d, *J* = 8.8 Hz, 2H), 7.59 – 7.44 (m, 2H), 7.37 (d, *J* = 8.3 Hz, 1H), 7.34 – 7.26 (m, 2H), 7.24 – 7.09 (m, 2H), 6.81 (dt, *J* = 15.4, 6.2 Hz, 1H), 6.40 (d, *J* = 15.4 Hz, 1H), 3.37 (d, *J* = 5.3 Hz, 2H), 2.39 (s, 6H).

THZ-P1-2-R: MS m/z 534.64 [M+H]^+. 1^H NMR (500 MHz, DMSO-*d*_6_) δ 12.06 (s, 1H), 10.32 (s, 1H), 10.23 (s, 1H), 9.49 (s, 1H), 8.76 (s, 1H), 8.27 (d, *J* = 3.0 Hz, 1H), 8.24 (d, *J* = 2.1 Hz, 1H), 8.12 (d, *J* = 7.7 Hz, 1H), 7.97 (d, *J* = 8.6 Hz, 2H), 7.75 (d, *J* = 8.8 Hz, 2H), 7.56 (d, *J* = 7.7 Hz, 1H), 7.52 (d, *J* = 8.1 Hz, 1H), 7.43 (d, *J* = 8.2 Hz, 1H), 7.41 – 7.34 (m, 2H), 7.26 (p, *J* = 7.0 Hz, 2H), 3.16 – 3.04 (m, 2H), 2.81 (d, *J* = 4.6 Hz, 6H), 2.47 (d, *J* = 7.1 Hz, 2H), 2.02 – 1.90 (m, 2H).

## Supplemental Information

**Figure S1.**
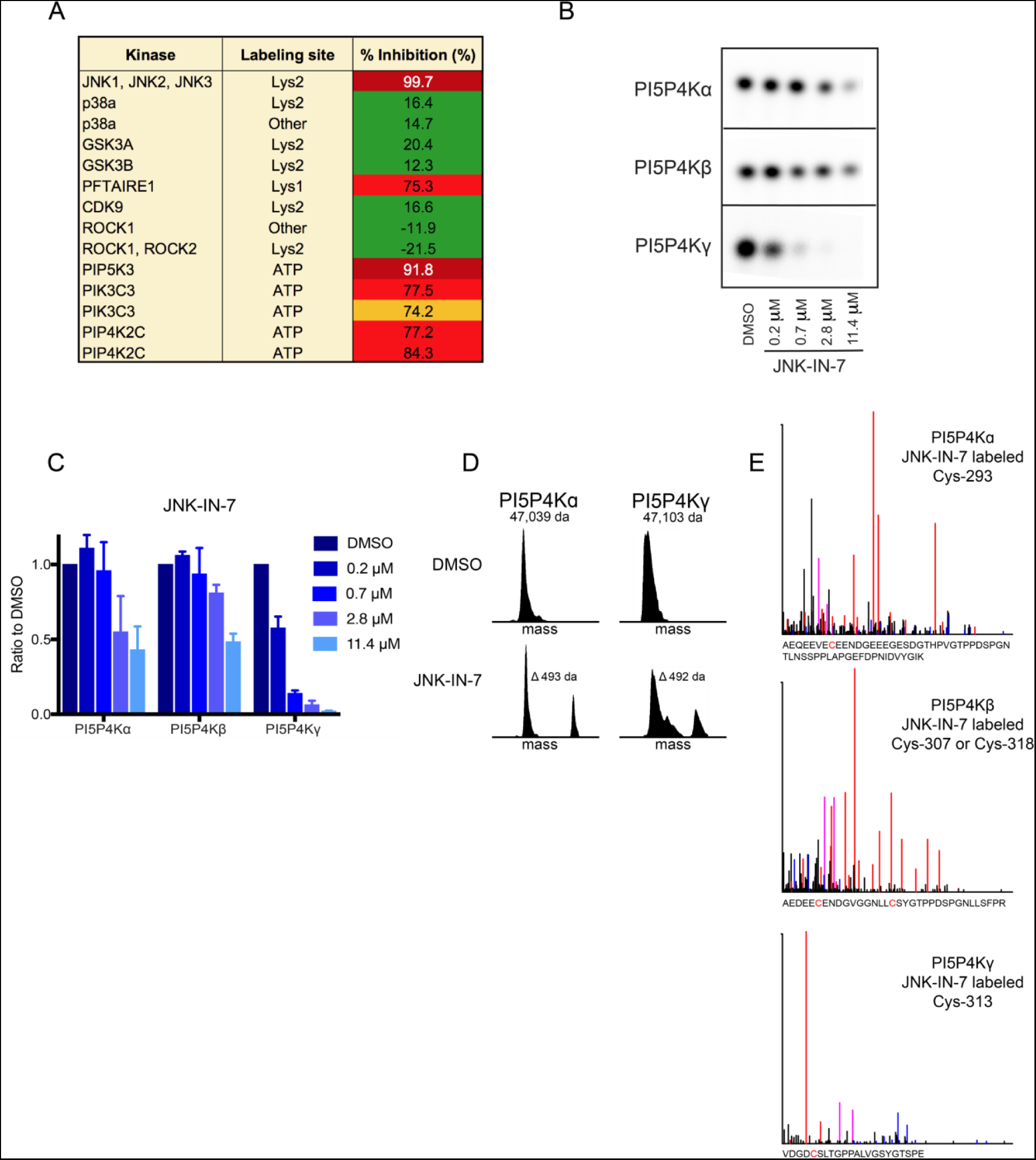
(A) list of selected targets, including PI5P4Kγ (PIP4K2C), from KiNativ profiling of JNK-IN-7 at 1 μM for 3 hours in A375 cells. A higher percent inhibition of kinase labeling by ATP-biotin is indicates stronger binding and inhibition of the target kinase. Full list is available in Zhang et al. (2012). (B) JNK-IN-7 inhibits kinase activity of all three PI5P4K isoforms in a radiometric TLC assay measuring radiolabeled PI-4,5-P_2_. JNK-IN-7 was incubated with purified protein for 30min, followed by a 10min kinase reaction, quenching, lipid extraction and TLC development. (C) Quantification of (B). Radiolabeled PI-4,5-P_2_ spots were imaged by autoradiography and quantified by densitometry. (D) Zero charge masses of recombinant PI5P4Kα and γ incubated with JNK-IN-7 for 2 hours at 37°C demonstrates covalent labeling of PI5P4K isoforms as determined by intact mass spectrometry. See Supplemental Intact MS Spectra for representative raw MS spectra used for charge deconvolution. (E) Subsequent protease digestion and tandem mass spectrometry confirms that THZ-P1-2 covalently labels cysteine residues on all three PI5P4K isoforms. The peptide for PI5P4Kβ was exclusively observed to be singly labeled at either Cys-307 or Cys-318. See Supplemental Intact MS Spectra for representative raw MS spectra used for charge deconvolution.

**Figure S2.**
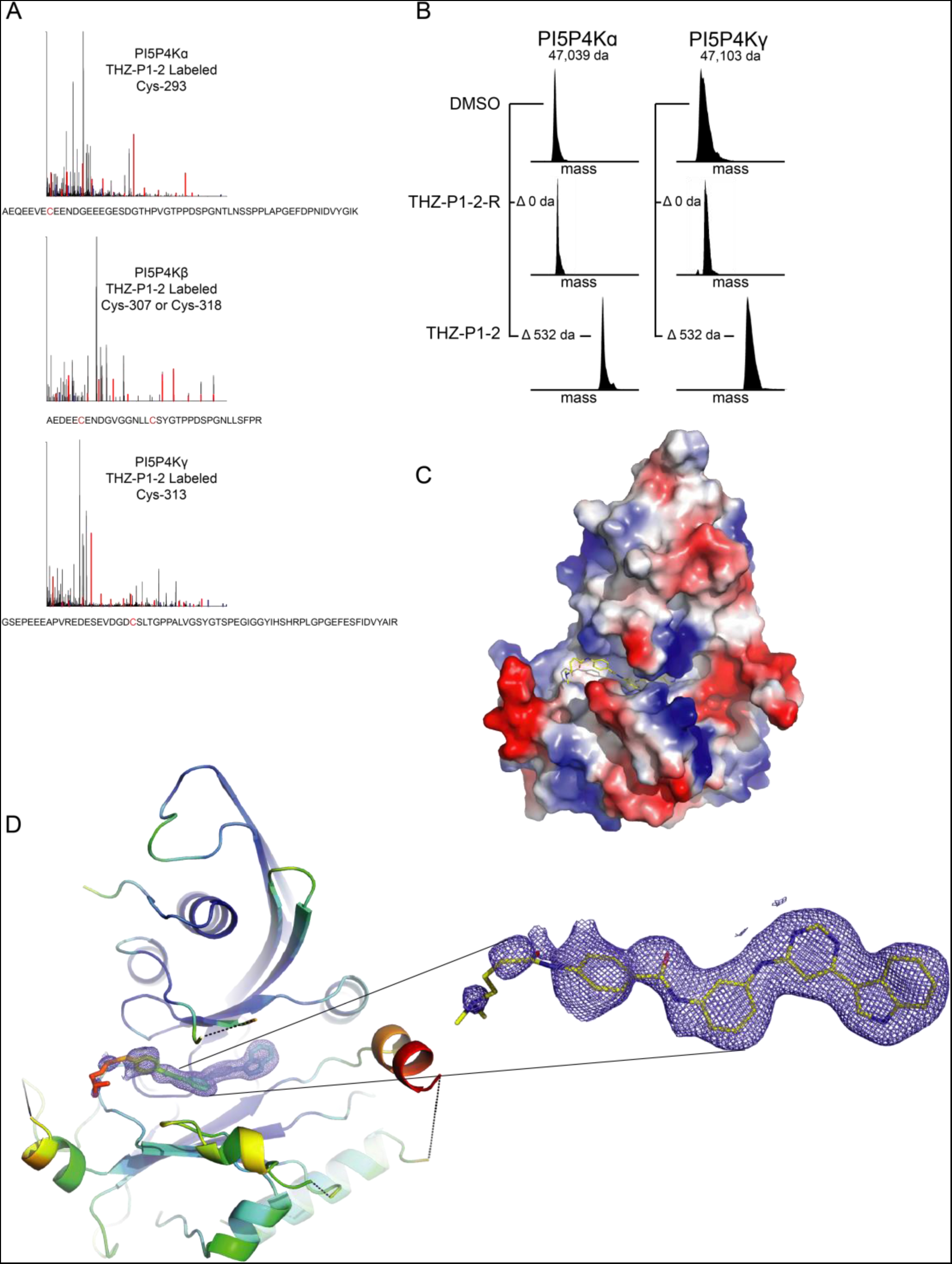
(A) Protease digestion and tandem mass spectrometry following intact mass labeling confirms that THZ-P1-2 covalently labels cysteine residues on all three PI5P4K isoforms. The peptide for PI5P4Kβ was exclusively observed to be singly labeled at either Cys-307 or Cys-318. Related to Fig. 2B. See Supplemental Intact MS Spectra for representative raw MS spectra used for charge deconvolution. (B) THZ-P1-2-R was found not to covalently label recombinant PI5P4Kα and γ when incubated with purified protein for 2 hours at 37°C. Mass labeling plot shown here is compared to mass shift observed with THZ-P1-2 from Fig. 2A. (C) Surface representation of co-crystal structure of PI5P4Kα in complex with THZ-P1-2. Related to Fig. 2C. (D) Zoom-in of ligand electron density within the active site of PI5P4Kα. See Table S1 for diffraction data collection and refinement statistics.

**Figure S3.**
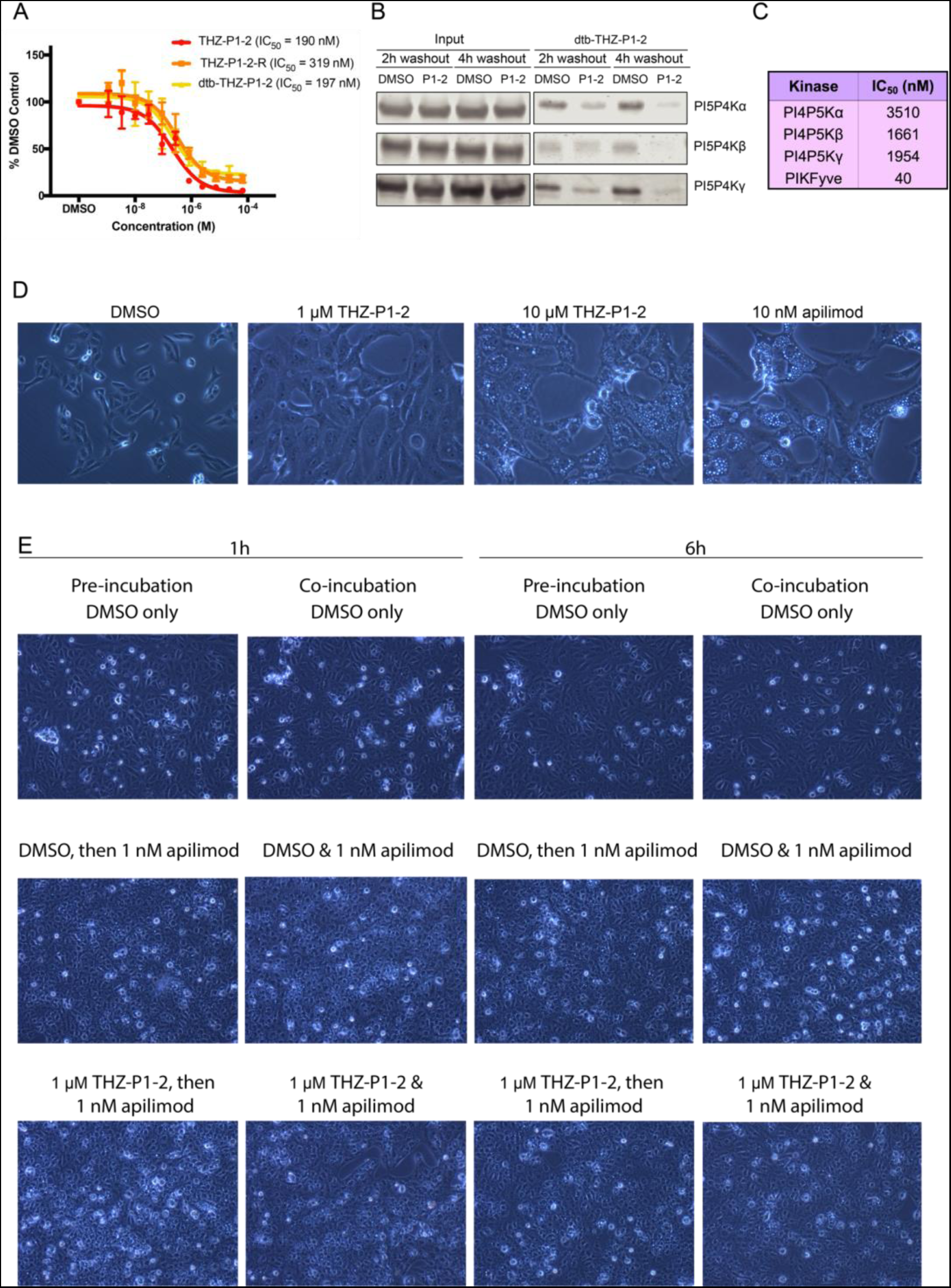
(A) THZ-P1-2-R and dtb-THZ-P1-2 were found to inhibit PI5P4Kα in an ADP-Glo assay, with a slight loss in potency with THZ-P1-2-R. Both compounds were incubated with PI5P4Ka for 15min before proceeding with a 1 hour reaction with ATP and development with ADP-Glo reagents. Plots are shown in comparison with THZ-P1-2 from Fig. 1B. (B) THZ-P1-2 engages PI5P4K isoforms at 2h and 4h timepoints, exhibiting prolonged engagement when a washout is performed. HEK293T cells were treated with DMSO or THZ-P1-2 at time, t=0 and cells were washed 2X with PBS at either 2 or 4h and harvested at the end of 6h. A streptavidin pulldown was then conducted in lysates after normalization of protein content with a BCA assay. (C) THZ-P1-2 inhibits the Type 1 PI4P5K kinases at a lower extent but shows potent inhibition on PIKfyve by ADP-Glo. *In vitro* kinase assays were performed by Carna Biosciences. (D) THZ-P1-2 causes vacuolar enlargement, characteristic of PIKfyve inhibition, at a concentration of 10 μ M but not 1 μ M, compared to PIKfyve inhibitor apilimod at 10 nM treatment. Vero cells were treated with DMSO or compound for 6h and imaged. (E) Incubation of Vero cells with 1 μM THZ-P1-2 was unable to impair the PIKfyve inhibitory phenotype of apilimod. Preincubation: Vero cells were incubated with DMSO/THZ-P1-2 for 1 or 6h, washed with PBS to completely remove compound, treated with DMSO/apilimod and imaged after 6h. Coincubation: Vero cells were incubated with DMSO/THZ-P1-2 for 1 or 6h, and then co-treated with DMSO/apilimod and imaged after 6h.

**Figure S4.**
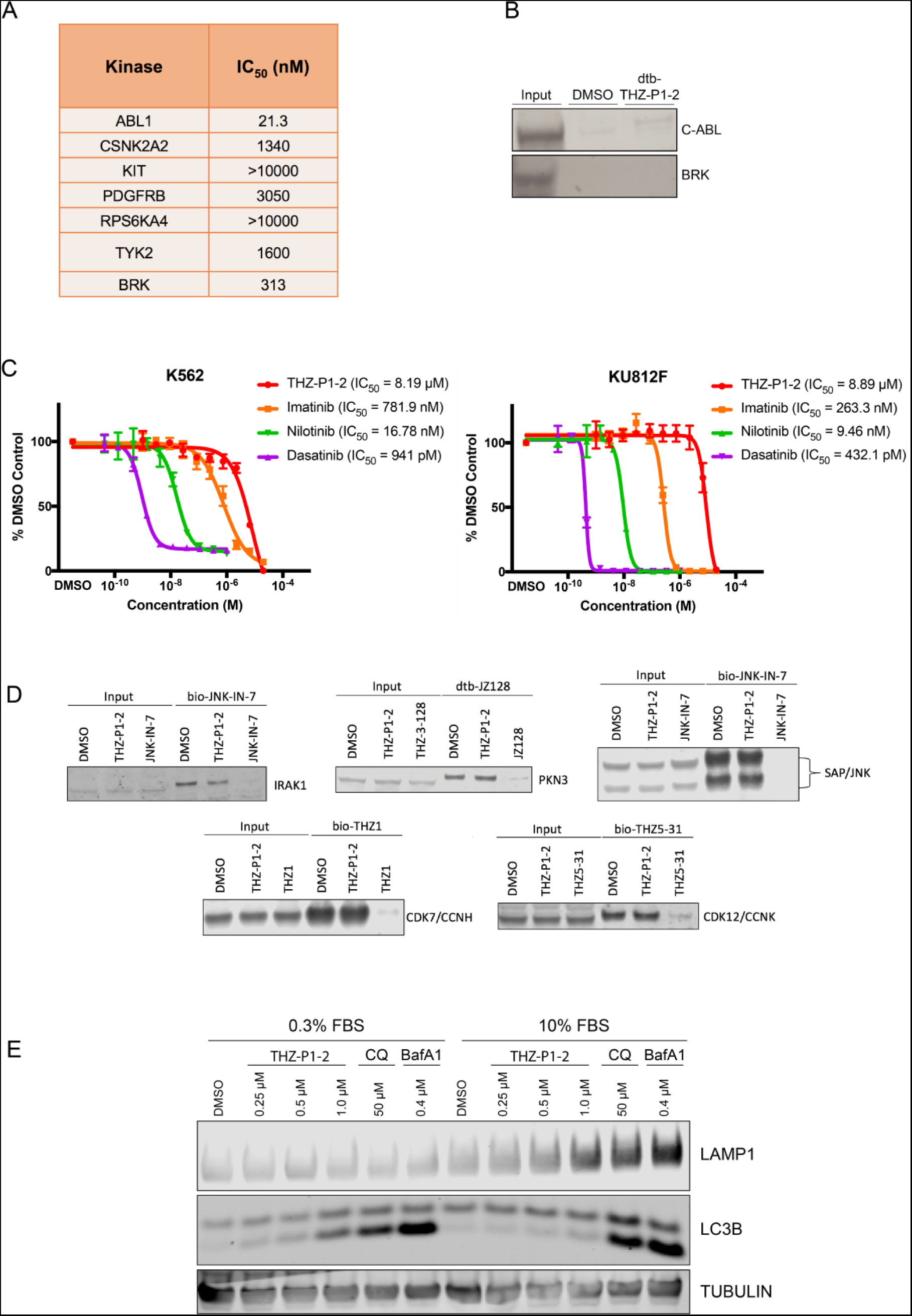
(A) THZ-P1-2 inhibits off-target kinases identified in the DiscoverX KINOMEScan panel to varying degrees. *In vitro* kinase assays were performed by Invitrogen using the Adapta assay. (B) A streptavidin pulldown in HEK293T lysate shows little to no engagement of C-ABL and BRK by dtb-THZ-P1-2. (C) THZ-P1-2 shows mild antiproliferative activity on BCR-ABL addicted cell lines. Two cell lines containing BCR-ABL translocations, K562 and KU812F, were treated with THZ-P1-2 or known BCR-ABL inhibitors imatinib, nilotinib and dasatinib for 72h and assayed using Cell-Titer Glo. (D) THZ-P1-2 does not bind to and engage off-targets such as JNK, IRAK1, PKN3, CDK7 and CDK12 despite originating from the same core scaffold as inhibitors of these targets. Competitive streptavidin pulldowns in HEK293T cells were conducted with DMSO, 1 μ M THZ-P1-2 and 1 μ M of each target’s corresponding inhibitor, followed by pulldown in lysate with 1 μ M of the corresponding biotinylated or desthiobiotinylated inhibitor. (E) THZ-P1-2 causes a slight increase in LAMP1 and LC3B protein levels, as observed with positive control compounds bafilomycin A1 and chloroquine. HeLa cells were cultured for 24h in DMEM media supplemented with either 0.3% (serum-starved conditions) or 10% FBS. Cells were treated with DMSO, varying concentrations of THZ-P1-2 and single doses of bafilomycin A1 and chloroquine, harvested after 24h, and analyzed by Western blot. Related to Fig. 4A.

**Table S1.** (Related to Figure 2C-D) Crystallization conditions, data collection and refinement statistics for co-crystal structure of complex THZ-P1-2 with PI5P4Kα

| Structure Name | PI5P4Kα/THZ-P1-2 |
| --- | --- |
| Ligand | THZ-P1-2 |
| RCSB accession code | 6OSP |
| **Data collection <sup>a</sup>** |  |
| Space group | P2<sub>1</sub> |
| Cell dimensions |  |
| *a*, *b*, *c* (Å) | 44.19 98.68 105.12 |
| *a*, *b*, *g* (°) | 90.00 93.66 90.00 |
| Resolution (Å) | 44.1-2.21 (2.29-2.21) <sup>b</sup> |
| *R*<sub>pim</sub> | 0.042 (0.525) |
| *I* / σ*I* | 13.44 (1.53) |
| Completeness (%) | 99.17 (99.44) |
| Redundancy | 3.7 (3.7) |
| **Structure solution** |  |
| PDB entries used for molecular replacement | 2YBX |
| **Refinement** |  |
| Resolution (Å) | 44.1-2.21 |
| No. reflections | 44717 |
| *R*<sub>work</sub> / *R*<sub>free</sub> | 0.1779/0.2254 |
| No. atoms | 5450 |
| Protein | 5102 |
| Ligand/ion | 92 |
| Water | 256 |
| *B*-factors |  |
| Protein | 53.08 |
| Ligand/ion | 70.65 |
| Water | 50.74 |
| R.m.s. deviations |  |
| Bond lengths (Å) | 0.012 |
| Bond angles (°) | 1.15 |
| **Ramachandran** |  |
| Preferred | 98.21% |
| Allowed | 1.63% |
| Not Allowed | 0.16% |

<sup>a</sup> A single crystal was used to collect data for each of the structures reported here.
<sup>b</sup> Values in parentheses are for highest-resolution shell.

## References

Adams, P. D., Afonine, P. V., Bunkóczi, G., Chen, V. B., Davis, I. W., Echols, N., … Zwart, P. H. (2010). PHENIX: A comprehensive Python-based system for macromolecular structure solution. Acta Crystallographica Section D: Biological Crystallography, 66(2), 213–221. https://doi.org/10.1107/S0907444909052925

Al-ramahi, I., Srinivas, S., Giridharan, P., Chen, Y., Patnaik, S., Safren, N., … Marugan, J. J. (2017). Inhibition of PIP4K g ameliorates the pathological effects of mutant huntingtin protein, 1–25.

Balla, T., Szentpetery, Z., & Kim, Y. J. (2009). Phosphoinositide Signaling: New Tools and Insights. Physiology, 24(4), 231–244. https://doi.org/10.1152/physiol.00014.2009

Braekeleer, E. D., Douet-Guilbert, N., Rowe, D., Bown, N., Morel, F., Berthou, C., … Braekeleer, M. D. (2011). ABL1 fusion genes in hematological malignancies: A review. European Journal of Haematology, 86(5), 361–371. doi:10.1111/j.1600-0609.2011.01586.x

Browne, C. M., Jiang, B., Ficarro, S. B., Doctor, Z. M., Johnson, J. L., Card, J. D., … Marto, J. A. (2018). A Chemoproteomic Strategy for Direct and Proteome-Wide Covalent Inhibitor Target-Site Identification. Journal of the American Chemical Society, 141(1), 191–203. doi:10.1021/jacs.8b07911

Bulley, S. J., Clarke, J. H., Droubi, A., Giudici, M. L., & Irvine, R. F. (2015). Exploring phosphatidylinositol 5-phosphate 4-kinase function. Advances in Biological Regulation, 57, 193–202. https://doi.org/10.1016/j.jbior.2014.09.007

Bulley, S. J., Droubi, A., Clarke, J. H., Anderson, K. E., Stephens, L. R., Hawkins, P. T., & Irvine, R. F. (2016). In B cells, phosphatidylinositol 5-phosphate 4-kinase-α synthesizes PI(4,5)P2 to impact mTORC2 and Akt signaling. Proceedings of the National Academy of Sciences of the United States of America, 113(38), 10571–10576. https://doi.org/10.1073/pnas.1522478113

Cai, X., Xu, Y., Cheung, A. K., Tomlinson, R. C., Alca, A., … Huang, Q. (2013). Article PIKfyve, a Class III PI Kinase, Is the Target of the Small Molecular IL-12 / IL-23 Inhibitor Apilimod and a Player in Toll-like Receptor Signaling, 912–921. https://doi.org/10.1016/j.chembiol.2013.05.010

Choy, C. H., Saffi, G., Gray, M. A., Wallace, C., Dayam, R. M., Ou, Z. A., … Botelho, R. J. (2018). Lysosome enlargement during inhibition of the lipid kinase PIKfyve proceeds through lysosome coalescence. https://doi.org/10.1242/jcs.213587

Clarke, J. H., Giudici, M., Burke, J. E., Williams, R. L., Maloney, D. J., Marugan, J., & Irvine, R. F. (2015). The function of phosphatidylinositol 5-phosphate 4-kinase γ (PI5P4K γ) explored using a specific inhibitor that targets the PI5P-binding site, 367, 359–367. https://doi.org/10.1042/BJ20141333

Cross, D. A., Ashton, S. E., Ghiorghiu, S., Eberlein, C., Nebhan, C. A., Spitzler, P. J., … Pao, W. (2014). AZD9291, an Irreversible EGFR TKI, Overcomes T790M-Mediated Resistance to EGFR Inhibitors in Lung Cancer. Cancer Discovery, 4(9), 1046–1061. doi:10.1158/2159-8290.cd-14-0337

Davis, M. I., Hunt, J. P., Herrgard, S., Ciceri, P., Wodicka, L. M., Pallares, G., … Zarrinkar, P. P. (2011). Comprehensive analysis of kinase inhibitor selectivity. Nature Biotechnology, 29(11), 1046–1051. doi:10.1038/nbt.1990

Davis, M. I., Sasaki, A. T., Shen, M., Emerling, B. M., Thorne, N., Michael, S., … Simeonov, A. (2013). A Homogeneous, High-Throughput Assay for Phosphatidylinositol 5-Phosphate 4-Kinase with a Novel, Rapid Substrate Preparation. PLoS ONE, 8(1). doi:10.1371/journal.pone.0054127

Emerling, B. M., Hurov, J. B., Poulogiannis, G., Tsukazawa, K. S., Choo-Wing, R., Wulf, G. M., … Cantley, L. C. (2013). XDepletion of a putatively druggable class of phosphatidylinositol kinases inhibits growth of p53-Null tumors. Cell, 155(4), 844–857. https://doi.org/10.1016/j.cell.2013.09.057

Emsley, P., & Cowtan, K. (2004). Coot: Model-building tools for molecular graphics. Acta Crystallographica Section D: Biological Crystallography, 60(12 I), 2126–2132. https://doi.org/10.1107/S0907444904019158

Fiume, R., Stijf-Bultsma, Y., Shah, Z. H., Keune, W. J., Jones, D. R., Jude, J. G., & Divecha, N. (2015). PIP4K and the role of nuclear phosphoinositides in tumour suppression. Biochimica et Biophysica Acta - Molecular and Cell Biology of Lipids, 1851(6), 898–910. https://doi.org/10.1016/j.bbalip.2015.02.014

Hasegawa, J., Strunk, B. S., & Weisman, L. S. (2017). PI5P and PI(3,5)P2: Minor, but Essential Phosphoinositides. Cell Structure and Function, 42(1), 49–60. doi:10.1247/csf.17003

Hu, A., Zhao, X., Tu, H., Xiao, T., Fu, T., Wang, Y., … Song, B. (2018). PIP4K2A regulates intracellular cholesterol transport through modulating PI(4,5)P2homeostasis. Journal of Lipid Research, 59(3), 507–514. doi:10.1194/jlr.m082149

Jude, J. G., Spencer, G. J., Huang, X., Somerville, T. D. D., Jones, D. R., Divecha, N., & Somervaille, T. C. P. (2014). A targeted knockdown screen of genes coding for phosphoinositide modulators identifies PIP4K2A as required for acute myeloid leukemia cell proliferation and survival. Oncogene, 34(10), 1–10. https://doi.org/10.1038/onc.2014.77

Kabsch, W. (2010). Integration, scaling, space-group assignment and post-refinement. Acta Crystallographica Section D Biological Crystallography, 66(2), 133–144. https://doi.org/10.1107/s0907444909047374

Karaman, M. W., Herrgard, S., Treiber, D. K., Gallant, P., Atteridge, C. E., Campbell, B. T., … Zarrinkar, P. P. (2008). A quantitative analysis of kinase inhibitor selectivity. Nature Biotechnology, 26(1), 127–132. doi:10.1038/nbt1358

Keune, W. J., Jones, D. R., & Divecha, N. (2013). PtdIns5P and Pin1 in oxidative stress signaling. Advances in Biological Regulation, 53(2), 179–189. https://doi.org/10.1016/j.jbior.2013.02.001

Keune, W. J., Jones, D. R., Bultsma, Y., Sommer, L., Zhou, X. Z., Lu, K. P., & Divecha, N. (2012). Regulation of phosphatidylinositol-5-phosphate signaling by Pin1 determines sensitivity to oxidative stress. Science Signaling, 5(252), 1–13. https://doi.org/10.1126/scisignal.2003223

Kimmelman, A. C., & White, E. (2017). Autophagy and Tumor Metabolism. Cell Metabolism, 25(5), 1037–1043. doi:10.1016/j.cmet.2017.04.004

Kitagawa, M., Liao, P., Lee, K. H., Wong, J., Shang, S. C., Minami, N., … Lee S. H. (n.d.). mitotic pathways leads to cancer-selective lethality. Nature Communications. https://doi.org/10.1038/s41467-017-02287-5

Kwiatkowski, N., Zhang, T., Rahl, P. B., Abraham, B. J., Reddy, J., Ficarro, S. B., … Gray, N. S. (2014). Targeting transcription regulation in cancer with a covalent CDK7 inhibitor. Nature, 511(7511), 616–20. https://doi.org/10.1038/nature13393

Lamia, K. A., Peroni, O. D., Kim, Y., Rameh, L. E., Kahn, B. B., & Cantley, L. C. (2004). Increased Insulin Sensitivity and Reduced Adiposity in Phosphatidylinositol 5-Phosphate 4-Kinase -/- Mice. Molecular and Cellular Biology, 24(11), 5080–5087. doi:10.1128/mcb.24.11.5080-5087.2004

Li, D., Ambrogio, L., Shimamura, T., Kubo, S., Takahashi, M., Chirieac, L. R., … Wong, K. (2008). BIBW2992, an irreversible EGFR/HER2 inhibitor highly effective in preclinical lung cancer models. Oncogene, 27(34), 4702–4711. doi:10.1038/onc.2008.109

Liu, Q., Sabnis, Y., Zhao, Z., Zhang, T., Buhrlage, S. J., Jones, L. H., & Gray, N. S. (2013). Developing irreversible inhibitors of the protein kinase cysteinome. Chemistry and Biology, 20(2), 146–159. https://doi.org/10.1016/j.chembiol.2012.12.006

Lundquist, M. R., Goncalves, M. D., Loughran, R. M., Possik, E., Vijayaraghavan, T., Yang, A., … Emerling, B. M. (2018). Phosphatidylinositol-5-Phosphate 4-Kinases Regulate Cellular Lipid Metabolism By Facilitating Autophagy. Molecular Cell, 70(3). doi:10.1016/j.molcel.2018.03.037

Ma, G., Duan, Y., Huang, X., Qian, C. X., Chee, Y., Mukai, S., … Lei, H. (2016). Prevention of Proliferative Vitreoretinopathy by Suppression of Phosphatidylinositol 5-Phosphate 4-Kinases. Investigative Opthalmology & Visual Science, 57(8), 3935. doi:10.1167/iovs.16-19405

Martelli A.M., Gilmour, R.S., Bertagnolo V., L.M. Neri, L. Manzoli, L. Cocco, Nuclear localization and signalling activity of phosphoinositidase C beta in Swiss 3T3 cells, Nature 358 (1992) 242–245.

McCoy, A. J., Grosse-Kunstleve, R. W., Adams, P. D., Winn, M. D., Storoni, L. C., & Read, R. J. (2007). Phaser crystallographic software. Journal of Applied Crystallography, 40(4), 658–674. https://doi.org/10.1107/s0021889807021206

Naviaux, R. K., Costanzi, E., Haas, M., & Verma, I. M. (1996). The pCL vector system: rapid production of helper-free, high-titer, recombinant retroviruses. Journal of Virology, 70(8), 5701–5. Retrieved from http://www.ncbi.nlm.nih.gov/pubmed/8764092%0A http://www.pubmedcentral.nih.gov/articlerender.fcgi?artid=PMC190538

N. Divecha, S.G. Rhee, A.J. Letcher, R.F. Irvine, Phosphoinositide signalling enzymes in rat liver nuclei: phosphoinositidase C isoform beta 1 is specifically, but not predominantly, located in the nucleus, Biochem. J. 289 (Pt 3) (1993) 617–620.

Pan, Z., Scheerens, H., Li, S., Schultz, B., Sprengeler, P., Burrill, L., … Palmer, J. (2007). Discovery of Selective Irreversible Inhibitors for Bruton’s Tyrosine Kinase. ChemMedChem, 2(1), 58–61. doi:10.1002/cmdc.200600221

Perera, R. M., Stoykova, S., Nicolay, B. N., Ross, K. N., Fitamant, J., Boukhali, M., … Bardeesy, N. (2015). Transcriptional control of autophagy-lysosome function drives pancreatic cancer metabolism. Nature, 524(7565), 361–365. https://doi.org/10.1038/nature14587

R. Fiume, W.J. Keune, I. Faenza, Y. Bultsma, G. Ramazzotti, D.R. Jones, A.M. Martelli, L. Somner, M.Y. Follo, N. Divecha, L. Cocco, Nuclear phosphoinositides: location, regulation and function, Subcell. Biochem. 59 (2012) 335–361.

Rabindran, S. K., Discafani, C. M., Rosfjord, E. C., Baxter, M., Floyd, M. B., Golas, J., … Wissner, A. (2004). Antitumor Activity of HKI-272, an Orally Active, Irreversible Inhibitor of the HER-2 Tyrosine Kinase. Cancer Research, 64(11), 3958–3965. doi:10.1158/0008-5472.can-03-2868

Rameh, L. E., Tolias, K. F., Duckworth, B. C., & Cantley, L. C. (1997). A new pathway for synthesis of phosphatidylinositol-4,5-bisphosphate. Nature, 390(6656), 192–196. doi:10.1038/36621

Rameh, L.E., and Cantley, L.C. (1999). The role of phosphoinositide 3-kinase lipid products in cell function. J. Biol. Chem. 274, 8347–8350.

Rosales-Rodríguez, B., Fernández-Ramírez, F., Núñez-Enríquez, J. C., Velázquez-Wong, A. C., Medina-Sansón, A., Jiménez-Hernández, E., … Rosas-Vargas, H. (2016). Copy Number Alterations Associated with Acute Lymphoblastic Leukemia in Mexican Children. A report from The Mexican Inter-Institutional Group for the identification of the causes of childhood leukemia. Archives of Medical Research, 47(8), 706–711. doi:10.1016/j.arcmed.2016.12.002

Settembre, C., Di Malta, C., Polito, V. A., Garcia Arencibia, M., Vetrini, F., Erdin, S., … Ballabio, A. (2011). TFEB links autophagy to lysosomal biogenesis. Science (New York, N.Y.), 332(6036), 1429–33. https://doi.org/10.1126/science.1204592

Sharma, G., Guardia, C. M., Roy, A., Vassilev, A., Saric, A., Griner, L. N., … DePamphilis, M. L. (2019). A Family of PIKFYVE Inhibitors with Therapeutic Potential Against Autophagy-Dependent Cancer Cells Disrupt Multiple Events in Lysosome Homeostasis. Autophagy, 0(0), 15548627.2019.1586257. https://doi.org/10.1080/15548627.2019.1586257

Shim, H., Wu, C., Ramsamooj, S., Bosch, K. N., Chen, Z., & Emerling, B. M. (2016). Deletion of the gene Pip4k2c, a novel phosphatidylinositol kinase, results in hyperactivation of the immune system, 113(27), 7596–7601. https://doi.org/10.1073/pnas.1600934113

Stijf-Bultsma, Y., Sommer, L., Tauber, M., Baalbaki, M., Giardoglou, P., Jones, D. R., … Divecha, N. (2015). The basal transcription complex component TAF3 transduces changes in nuclear phosphoinositides into transcriptional output. Molecular Cell, 58(3), 453–467. https://doi.org/10.1016/j.molcel.2015.03.009

Sumita, K., Lo, Y. H., Takeuchi, K., Senda, M., Kofuji, S., Ikeda, Y., … Sasaki, A. T. (2016). The Lipid Kinase PI5P4Kβ Is an Intracellular GTP Sensor for Metabolism and Tumorigenesis. Molecular Cell, 61(2), 187–198. https://doi.org/10.1016/j.molcel.2015.12.011

Urayama, K. Y., Takagi, M., Kawaguchi, T., Matsuo, K., Tanaka, Y., Ayukawa, Y., … Manabe, A. (2018). Regional evaluation of childhood acute lymphoblastic leukemia genetic susceptibility loci among Japanese. Scientific Reports, 8(1). doi:10.1038/s41598-017-19127-7

Vicinanza, M., Korolchuk, V. I., Ashkenazi, A., Puri, C., Menzies, F. M., Clarke, J. H., & Rubinsztein, D. C. (2015). PI(5)P regulates autophagosome biogenesis. Molecular Cell, 57(2), 219–234. https://doi.org/10.1016/j.molcel.2014.12.007

Voss, M. D., Czechtizky, W., Li, Z., Rudolph, C., Petry, S., Brummerhop, H., … Schaefer, H. (2014). Discovery and pharmacological characterization of a novel small molecule inhibitor of phosphatidylinositol-5-phosphate 4-kinase, type II, beta. Biochemical and Biophysical Research Communications, 449(3), 327–331. doi:10.1016/j.bbrc.2014.05.024

Zhang, T., Inesta-Vaquera, F., Niepel, M., Zhang, J., Ficarro, S. B., MacHleidt, T., … Gray, N. S. (2012). Discovery of potent and selective covalent inhibitors of JNK. Chemistry and Biology, 19(1), 140–154. https://doi.org/10.1016/j.chembiol.2011.11.010

Zhang, T., Kwiatkowski, N., Olson, C. M., Dixon-clarke, S. E., Abraham, B. J., Greifenberg, A. K., … Gray, N. S. (2016). Covalent targeting of remote cysteine residues to develop CDK12 and CDK13 inhibitors. Nature Chemical Biology, 12(10), 876–884. https://doi.org/10.1038/nchembio.2166

Zheng, L., & Conner, S. D. (2018). PI5P4Kγ functions in DTX1-mediated Notch signaling. Proceedings of the National Academy of Sciences, 115(9). doi:10.1073/pnas.1712142115

